# Acetylated Chromatin Domains Link Chromosomal Organization to Cell- and Circuit-level Dysfunction in Schizophrenia and Bipolar Disorder

**DOI:** 10.1101/2021.06.02.446728

**Authors:** Kiran Girdhar, Gabriel E. Hoffman, Jaroslav Bendl, Samir Rahman, Pengfei Dong, Will Liao, Leanne Brown, Olivia Devillers, Bibi S. Kassim, Jennifer R Wiseman, Royce Park, Elizabeth Zharovsky, Rivky Jacobov, Elie Flatow, Alexey Kozlenkov, Thomas Gilgenast, Jessica S. Johnson, Lizette Couto, Mette A. Peters, Jennifer E Phillips-Cremins, Chang-Gyu Hahn, Raquel E. Gur, Carol A. Tamminga, David A. Lewis, Vahram Haroutunian, Psychencode Consortium, Stella Dracheva, Barbara K. Lipska, Stefano Marenco, Marija Kundakovic, John F. Fullard, Yan Jiang, Panos Roussos, Schahram Akbarian

**Author notes:** List of PsychENCODE consortium members is provided in the appendix (Psychencode Consortium). Equally contributing senior authors (Panos Roussos, Schahram Akbarian).

## Abstract

To explore modular organization of chromosomes in schizophrenia (SCZ) and bipolar disorder (BD), we applied ‘population-scale’ correlational structuring of 739 histone H3-lysine 27 acetylation and H3-lysine 4 trimethylation profiles, generated from the prefrontal cortex (PFC) of 568 cases and controls. Neuronal histone acetylomes and methylomes assembled as thousands of cis-regulatory domains (CRDs), revealing fine-grained, kilo-to megabase scale chromatin organization at higher resolution but firmly integrated into Hi-C chromosomal conformations. Large clusters of domains that were hyperacetylated in disease shared spatial positioning within the nucleus, predominantly regulating PFC projection neuron function and excitatory neurotransmission. Hypoacetylated domains were linked to inhibitory interneuron- and myelination-relevant genes. Chromosomal modular architecture is affected in SCZ and BD, with hyperacetylated domains showing unexpectedly strong convergences defined by cell type, nuclear topography, genetic risk, and active chromatin state across a wide developmental window.

## Introduction

Regulation of nucleosomal histone modifications, including active chromatin-associated lysine acetylation and methylation, is one of the top scoring biological pathways in genome-wide association studies (GWAS) of schizophrenia (SCZ) and related co-heritable traits, including bipolar disorder (BD)^1^. In addition, SCZ and BD risk loci are enriched for active neuronal promoters and enhancers in the adult human frontal lobe ^2–5^. Furthermore, rare mutations in a subset of histone modifying enzymes, regulating methylation and acetylation at sites of neuronal gene expression, show high disease penetrance^6,7^. Since representative genome-scale histone modification studies in brain disease are lacking, it remains unknown whether changes in histone acetylation and methylation landscapes affect the general population of subjects diagnosed with SCZ and BD^8^.

In conventional brain epigenomic maps, histone marks associated with transcription, including trimethyl-H3K4 (H3K4me3) and acetyl-H3K27 (H3K27ac), are enriched at promoters and enhancers and appear as isolated ‘peaks’. These peaks are mostly confined to short nucleosomal arrays, typically covering an average of 3.6-3.8 kb ^2,9^. Furthermore, only a very small portion of peaks show some degree of confluence by merging into super-enhancers important for cell-specific gene expression programs^10^. However, histone modification landscapes are tightly linked to chromatin structures, defined by local chromosomal conformations, including the megabase-scaling ‘self-folded’ topologically associating domains (TADs) and other features of chromosomal architecture and 3D genome organization ^11^. Whether or not such types of acetyl- and methyl-histone defined higher order chromatin structures exist in the human brain, including their cell type-specific regulation, potential alterations in disease and implications for the underlying genetic risk architecture, remain unknown.

Here, we generated 739 (N_H3K4me3_=230, N_H3K27ac_= 260 from neurons and N_H3K27ac_= 249 from bulk tissue) ChIP-seq (chromatin immunoprecipitation followed by deep sequencing) libraries from SCZ, BD and control brains from four independent brain collections. Using population-scale correlational analysis and cell type-specific chromosomal conformation mapping from adult prefrontal cortex (PFC), we define acetylation and methylation landscapes of neuronal chromatin by the coordinated regulation of sequentially arranged histone peaks constrained by local chromosomal conformations and nuclear topographies. We report widespread disease-associated alterations affecting the neuronal H3K27ac acetylome, but not the H3K4me3 methylome. On a genome-wide scale, hundreds of kilo- to megabase-scale chromosomal domains showed evidence for dysregulated acetylation, with strikingly converging alignments by genetic risk, cell type, developmental function, nuclear topography, and active vs. repressive chromosomal environments. Our findings, which were surprisingly consistent across multiple sets of case control studies, each with a different demographic distribution, strongly hint at a higher order chromatin-defined brain pathology representative of the broader population of subjects diagnosed with SCZ and BD.

## Results

### Acetyl-histone peaks show disease specific dysregulation

We first generated genome-wide maps of histone modifications, H3K4me3 and H3K27ac in neuronal (NeuN+) nuclei isolated from dorsolateral PFC via fluorescence activated nuclear sorting (FANS). We generated 490 ChIP-seq libraries (approximately 3-5×10^5^ NeuN^+^ nuclei/PFC sample) from 321 demographically matched SCZ and non-psychiatric control brains, hereafter referred to as study-1. Donor specimens are part of the CommonMind Consortium collection ^12–14^, provided by three brain collections (The Mount Sinai/JJ Peters VA Medical Center NIH Brain and Tissue Repository (MSSM), the University of Pittsburgh Brain Tissue Donation Program (PITT) and the UPenn Center for Neurodegenerative Disease Research (PENN)). We generated an additional set of H3K27ac ChIP-seq libraries (N=249) prepared from unsorted nuclei extracted from bulk dorsolateral PFC tissue of SCZ, BD and control brains. These specimens, hereafter referred to as study-2, were contributed by the Human Brain Collection Core (HBCC) at the NIMH (***Figure 1A, Table 1, Table S1A-B***).

**Figure 1:**
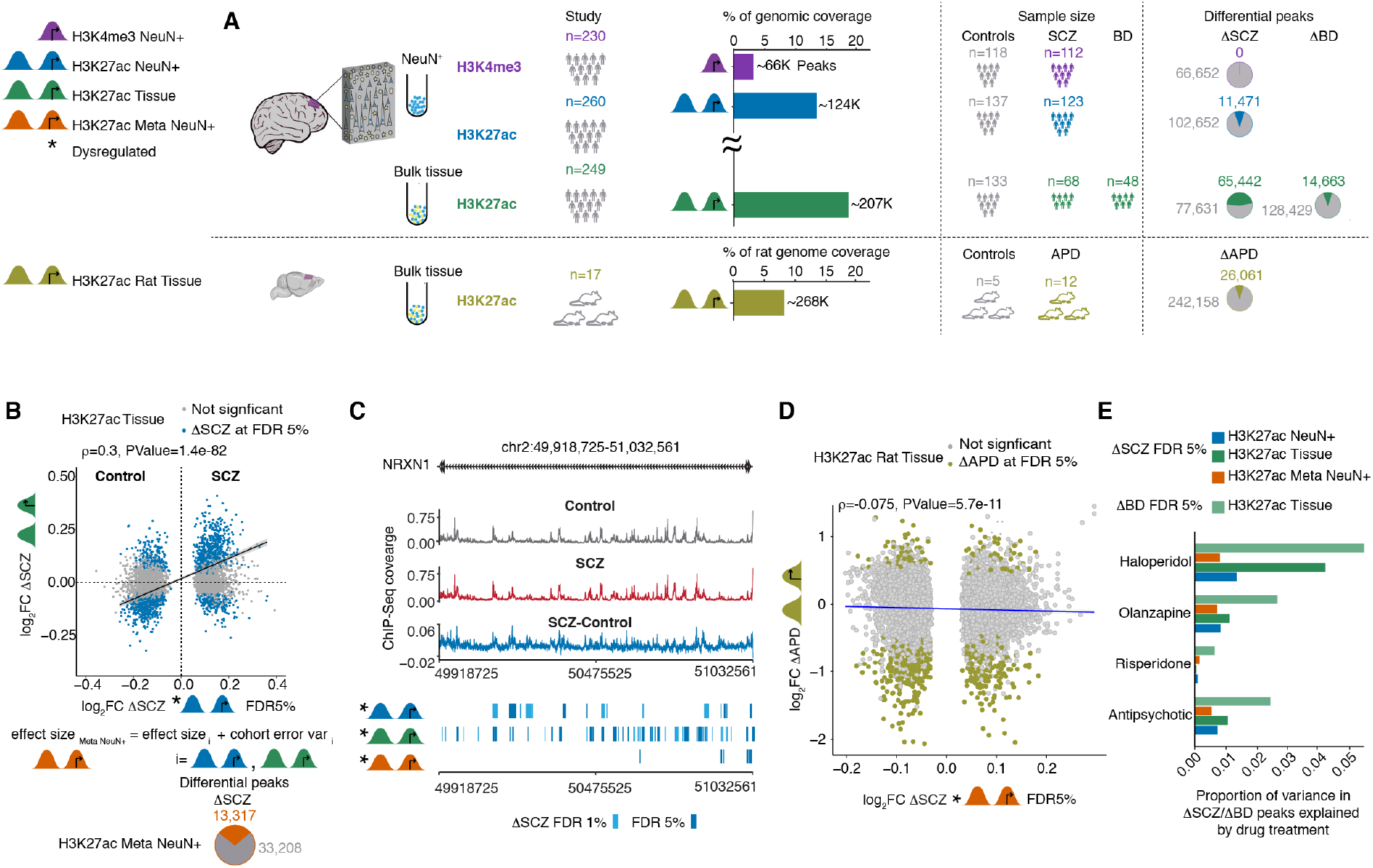
Histone peak profiling in 739 ChIP-Seq datasets from two studies consisting of SCZ, BD and control subjects. **(A)** (left) Datasets and studies; study-1, FANS isolated PFC NeuN^+^ nuclei: H3K4me3 (purple), H3K27ac (blue). study-2, total (non-sorted) tissue PFC nuclei: H3K27ac (green). Meta-analysis study-1 and -2, H3K27ac (orange). Rat study, Frontal cortex total (non-sorted) tissue nuclei H3K27ac (brown-green). (middle) Bar plot, genomic coverage (%) of each ChIP-Seq dataset. Numbers of subjects and animals as indicated. Rat APD treatments (17 animals), N=4 olanzapine, N=3 haloperidol, N=5 risperidone, N=5 vehicle controls. (Right) Pie charts (colored sectors marking significantly different peaks) showing numbers and proportion of differentially regulated histone peaks in (top 3 rows) case control study comparison, or (bottom row) antipsychotic treatment versus control. **(B)** Correlation of effect sizes of H3K27ac NeuN+ peaks altered in SCZ study-1, compared with effect sizes for corresponding peaks in SCZ study-2 subjects with bulk PFC Tissue as input. Blue dots mark H3K27ac NeuN+ at FDR < 5%. Meta analysis of effect sizes of H3K27ac Meta NeuN+ and H3K27ac Tissue using a fixed effect model generated H3K27ac Meta NeuN+ peakset. The pie chart shows differential SCZ specific histone modified regions of H3K27ac Meta NeuN+ at FDR < 5%. **(C)** Visualization of over 1Mb region covering NRXN1 gene to show ChIP-sSeq coverage of H3K27ac NeuN+ separately from control individuals (gray color, top panel), and subjects with SCZ individuals (red color, second panel) and difference between the coverage (blue color, third panel) in H3K27ac Meta NeuN+ (orange), H3K27ac Tissue (green), H3K27ac NeuN+ (blue). **(D)** Correlation of effect sizes of H3K27ac Meta NeuN+ peaks altered in SCZ, compared with effect sizes for corresponding peaks in antipsychotics treated rats with PFC H3K27ac Tissue as input. Olive green dots mark Rat H3K27ac Tissue peaks altered due to antipsychotic treatment at FDR < 5%. **(E)** Proportion of variance in SCZ dysregulated peaks in study-1 H3K27ac NeuN+, study-1 H3K27ac Meta NeuN+, study-2 H3K27ac Tissue and BD dysregulated peaks in study-2 H3K27ac Tissue explained by antipsychotics (olanzapine, haloperidol, and risperidone) as indicated.

**Table 1:** Summary of metadata of samples in study-1 and study-2

| Study | Marker | PFC cell type | Brain bank | Diagnosis | Gender | Age | Ethnicity |
| --- | --- | --- | --- | --- | --- | --- | --- |
| Study-1 | H3K4me3 | NeuN+ | MSSM<br>PENN<br>PITT | 112 SCZ<br>118 Ctrl | 92F/138M | 66.33±18.42 | 183 CAU, 34 AA, 2 EA,<br>10 HISP, 1 Multiracial |
|  | H3K27ac |  |  | 123 SCZ<br>137 Ctrl | 98F/162M | 67.32±18.02 | 203 CAU, 38 AA, 2 EA,<br>16 HISP, 1 Multiracial |
| Study-2 | H3K27ac | Bulk tissue | HBCC | 68 SCZ, 48 BD<br>133 Ctrl | 88F/161M | 42.27±17.59 | 121 CAU, 117 AA, 6 EA,<br>5 HISP |

SCZ: Schizophrenia, BD: Bipolar Disorder, Ctrl: Control CAU: Caucasian, AA: African-American, EA: East Asian, HISP: Hispanic

After applying quality control steps and regressing out various technical factors related to tissue processing and sequencing (***Methods, Figure S1A-B***), we obtained three sets of normalized histone peak activity matrices (64,254 peaks × 230 H3K4me3 NeuN+, 114,136 peaks × 260 H3K27ac NeuN+ from study-1 and 143,074 peaks × 249 H3K27ac Tissue from study-2; ***Tables S2A-C***). Of note, for each study-2 sample, cell type heterogeneity of the bulk tissue was adjusted by estimating the proportion of neurons using cell type specific ChIP-seq data generated in an independent set of donors^2^ (***Figure S1B***). As expected, the narrow-defined H3K4me3 peaks had lower genomic coverage (3.1%) compared to the broad H3K27ac marker (12.8% in neurons and 17.1% in bulk tissue; ***Figure 1A***). More than 60% of H3K4me3 and 75% of H3K27ac peaks were distributed among distal intergenic, exonic, intronic and UTR elements (***Figure S1C***). Importantly, each dataset showed high concordance, measured as Jaccard similarity coefficients (∼0.7), to previously generated H3K4me3 and H3K27ac datasets from PFC of independent sets of brain samples not included in the present study (***Figure S1D***) ^2^.

We first explored histone ‘peak’-based disease-associated epigenomic aberrations in the promoter associated H3K4me3 regions in the PFC of SCZ cases vs controls. Surprisingly, none of the 64,254 peaks (***Table S3A***) survived multiple testing correction after differential (cases vs controls) analysis, indicating that this methylation mark is not consistently affected in SCZ PFC. Next, we evaluated SCZ specific epigenomic aberrations in the promoter and enhancer associated H3K27ac marked regions of the PFC. Altogether, 11,471 of the 114,136 H3K27ac NeuN+ peaks were dysregulated in SCZ study-1, at FDR 5% (***Table S3B***), and an even larger proportion of H3K27ac Tissue was significantly affected in SCZ study-2 (***Figure 1A, Table S3C***). However, there was a robust correlation between case-control effect size of SCZ H3K27ac NeuN+ at FDR 5 % and case-control effect size of SCZ H3K27ac Tissue (Spearman’s ρ=0.3, P= 1.4 ×10^−82^) (***Methods, Figure 1B, Figure S2A***). Having shown that histone acetylation changes in SCZ PFC are broadly reproducible across independent brain collections and studies (***Table S1***), we next combined the differential histone peak effects sizes and p-values from study-1 H3K27ac NeuN+ and study-2 H3K27ac Tissue datasets (***Methods***), yielding a consensus set of 46,345 H3K27ac Meta NeuN+ peaks, out of which 13,137 peaks were dysregulated in SCZ, at FDR 5% (***Figure 1B, Figure S2B, Table S3D***). We applied a similar differential analysis workflow to determine BD specific epigenomic aberrations in study-2 H3K27ac Tissue marked regions. Altogether, 14,663 out of 143,074, peaks were identified as dysregulated, at FDR 5% (***Table S3E)***. Finally, there was a robust correlation of BD- and SCZ-sensitive peaks across the studies (H3K27ac NeuN+, H3K27ac Tissue) and within the same cohort, or SCZ and BD from study-2 H3K27ac Tissue, (***Figure S3A*)**, suggesting shared epigenomic dysfunction in these two common types of psychiatric disorders. Genes set enrichment analysis of dysregulated peaks were remarkably consistent in pattern across our SCZ and BD cohorts, and neuronal (including neurodevelopmental) signaling and synaptic plasticity pathways ranked top among gene ontologies of disease-sensitive H3K27ac peaks (***Figure S4*)**. A representative example of dysregulated H3K27ac peak-based alterations in our SCZ PFC datasets (study-1, study-2, meta) is shown in ***Figure 1C***, with the well studied 1.1Mb wide Neurexin 1 (*NRXN1)* cell adhesion gene and neurodevelopmental susceptibility locus^15^.

Because the majority of our diseased brains were exposed to antipsychotic drugs (APD) prior to death (***Table S1A-B***), we wanted to assess the potential impact of medication. To this end, we prepared H3K27ac ChIP-seq from N=17 (plus an additional N=5 deeply sequenced nucleosomal ‘input’ preparations) adult rat rostro-medial frontal cortex continuously exposed to APD in the drinking water for a period of 4.5 months^16,17^ (***Figure S5A-C, Table S5A-C***). We identified 26,061 peaks sensitive to APD at FDR 5%, with very high (0.9 - 0.99) pairwise correlations across the three APD groups, including D_2_-like antagonist haloperidol (2 mg/kg/day, N=3 animals), the atypical (dopaminergic/serotonergic) APD olanzapine (4 mg/kg/day, N=4 animals) and risperidone (6mg/kg/day, N=5 animals), compared to water controls (N=5 animals). We conclude that APDs share an epigenomic response profile (***Figure S5D-F***). We identified, by Hg38 liftover, APD-sensitive (rat) peaks dysregulated in our schizophrenia cases. However, given the very weak correlations (ρ = -0.015 (SCZ H3K27ac Meta NeuN+), ρ = -0.087 (SCZ H3K27ac NeuN+), and ρ = -0.1 (SCZ H3K27ac Tissue) and ρ = -0.16 (BD H3K27ac Tissue) (***Figure 1D, Figure S5G-I***), with only a very small portion of variance, (0.0005-0.0056) in differential SCZ and BD peak profiles explained by medication, we conclude that the vast majority of H3K27ac peak alterations in diseased PFC are not explained by APD exposure ***(Figure 1E)***. Instead, many disease-sensitive peaks including the *BDNF* neurotrophin, the *DLGAP3* glutamate receptor anchoring protein and other key regulators of synaptic signaling showed inverse changes in diseased human PFC and APD-exposed rat cortex (***Table S5C***).

### Enhancer peaks are enriched for disease signatures and common risk variants

We hypothesized that epigenomic alterations in diseased PFC predominantly occur at the site of enhancers, because H3K4me3, which is primarily associated with promoters ^18^, was not affected in our cases, as opposed to the widespread changes observed for the enhancer-associated mark, H3K27ac. To this end, we divided the H3K27ac peak-sets into two categories: 1) promoters (± 3Kb from TSS) and 2) enhancers (peaks other than promoters). Indeed, SCZ- and BD-specific H3K27ac-peak alterations showed strong enrichment for enhancer-specific elements, with significant depletion for promoter specific elements from following studies 1) SCZ H3K27ac NeuN+, 2) SCZ H3K27ac Tissue and 3) BD H3K27ac Tissue (***Figure S6A***). Next, we tested the association of common genetic variants from a diverse set of psychiatric and behavioral traits in disease altered enhancer elements, promoter elements and combined (promoters + enhancers) elements in the above mentioned three studies. Enrichment analysis for SCZ risk variants using stratified LD score regression ^19^, showed that the normalized heritability coefficients were particularly strong for enhancer associated, SCZ specific H3K27ac peaks in comparison to promoter and combined (promoter + enhancer) peaksets (***Figure 2A***). These changes were highly specific because non-psychiatric traits, such as height, or medical conditions including but not limited to autoimmune and cardiac disease completely lacked association with the disease-associated PFC peaks of the present study (***Figure S6B, Table S4A***). BD specific H3K27ac peaks showed significant heritability coefficients for SCZ but not BD risk variants when stratified by promoters and enhancers (***Figure S7***), which could reflect underpowered Ld score enrichment test as the number of GWAS loci in BD are less than SCZ.

**Figure 2:**
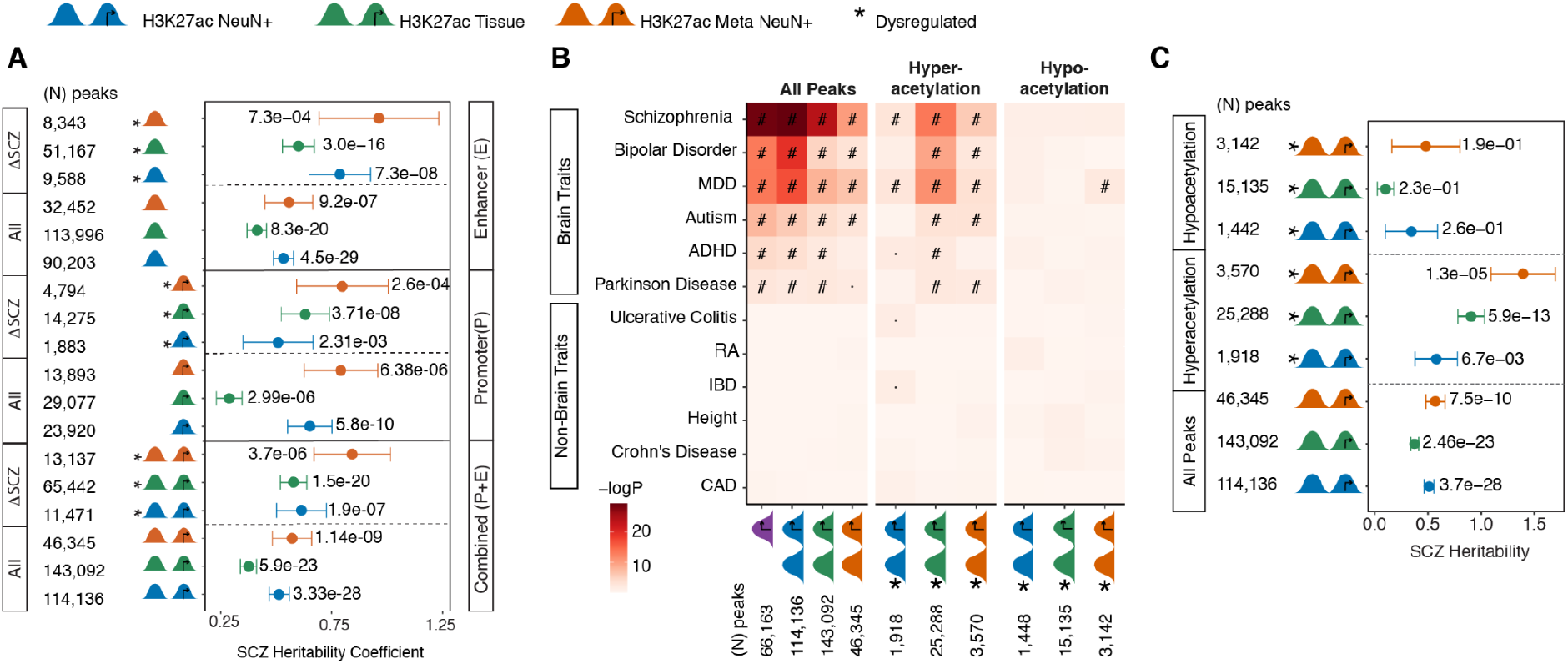
Enrichment of common SCZ risk variants in dysregulated peaks in NeuN+ and bulk tissue. **(A)** SCZ heritability coefficients of genetic variants overlapping histone peaks from study-1 H3K27ac NeuN+, study-2 H3K27ac Tissue and H3K27ac Meta NeuN+. The heritability coefficient plot in A) is stratified by 1) “All”: all histone peaks and 2) “ΔSCZ”: dysregulated peaks categorized as Combined (P+E) which is promoters + enhancers, Promoters (P) :peaks thats are within ±3Kb from TSS) and Enhancers (E): peaks other than promoters peaks. The overlap of peaks with genetic variants was assessed using LD score regression. “#”: Significant for enrichment in LD score regression after FDR correction of multiple testing across all tests in the plot (Benjamini & Hochberg); “·”: Nominally significant for enrichment. **(B)** Heatmap of enrichment *P*-values of brain and non-brain related GWAS traits and **(C)** heritability coefficients of common genetic variants in SCZ overlapping histone peaks from H3K27ac NeuN+, H3K27ac Tissue and H3K27ac Meta NeuN+. The heatmap in B) and heritability coefficient plot in C) are stratified by 1) “All peaks”: genome wide histone peaks, 2) “hyperacetylated”: dysregulated peaks in SCZ vs controls comparison with log_2_FC > 0 & adj.P.value <.01 and “hypoacetylated”: dysregulated peaks in SCZ vs controls comparison with log_2_FC>0 & adj.P.value <.01. The overlap of peaks with genetic variants was assessed using LD score regression. “#”: Significant for enrichment in LD score regression after FDR correction of multiple testing across all tests in the plot (Benjamini & Hochberg); “·”: Nominally significant for enrichment.

### Hyper- but not hypoacetylated peaks are linked to SCZ risk variants

To better understand these disease-associated aberrations in PFC H3K27ac peaks, we stratified altered peaks into hypoacetylation (log_2_ fold change <0) and hyperacetylation (log_2_ fold change >0). Interestingly, in SCZ PFC, the association between genetic risk and dysregulated acetylation was overwhelmingly driven by the group of hyper-but not hypo-acetylated peaks in all three of our SCZ case control comparisons (H3K27ac NeuN+, H3K27ac Tissue and H3K27ac Meta NeuN+) (***Figure 2B***). A strikingly similar pattern of alterations emerged in our BD focused analysis (study-2) (***Figure S7, Table S3E)***. In contrast, hypoacetylated peaks in SCZ and BD were void of any association with genetic risk (***Figure 2A-B, Figure S7***).

### Reconstruction of modular, ‘3D’ chromosomal architecture by H3K4me3 and H3K27ac correlational patterning

After identifying alterations in the activity of PFC histone peaks, we investigated the impact of disease on structural organization of PFC chromatin by characterizing the modular architecture of coordinated histone peaks in brain epigenome. We hypothesized that coordinated structure of chromatin peaks could be particularly important in the context of disease. This is a plausible hypothesis given the examples from peripheral cells and tissues for which coordinated regulation of multiple cis-regulatory elements, often sequentially organized along the linear genome, has been reported ^20^. Indeed, ‘population-scale’ interindividual correlations between histone peaks was performed on hundreds of lymphoblastoid and fibroblast cultures and has uncovered coordinated regulation, or ‘cis-regulatory domains’ (CRDs). CRDs span spatial clustering of chromatin peaks that extends across 10^4^-10^6^ base pairs of linear genome sequence and integrate into local chromosomal conformation landscapes ^21,22^. Similar approaches have been successfully applied to open chromatin regions in postmortem brains from donors with Alzheimer’s disease ^23^.

Here, we developed a systematic workflow (***Methods, Figure S8***) by combining the previously developed software *decorate* ^24^ *with additional steps of statistical analyses to identify CRDs on our population-scale H3K27ac and H3K4me3 datasets encompassing 739 PFC ChIP-seq libraries. The pipeline applied adjacency constrained hierarchical clustering* ^24,25^, *across each of our three ChIP-seq datasets (H3K4me3 NeuN+, H3K27ac NeuN+, H3K27ac Tissue) to identify sequentially aligned clusters of peaks as a strongly correlated structure (****Methods, Figure S9A-B***). Altogether, 39% (H3K4me3 NeuN+), 65% (H3K27ac NeuN+) and 69% (H3K27ac Tissue) of peaks assembled into 2,721, 6,389 and 8,247 CRDs respectively (***Figure 3A, Table S6A-C***), with H3K27ac CRDs encompassing on average ∼11.5 histone peaks, whereas H3K4me3 CRDs averaged at 9.1 histone peaks (***Figure S9C***).

**Figure 3:**
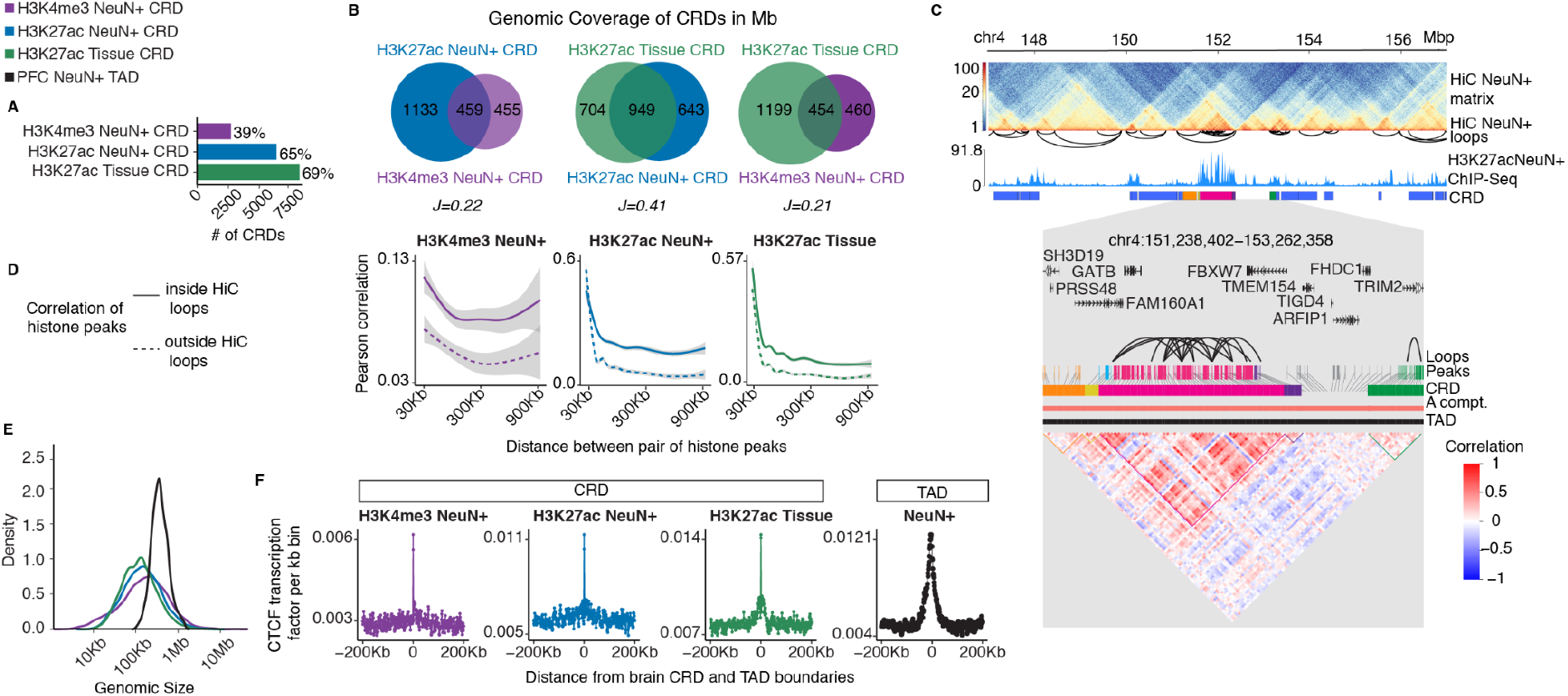
PFC histone CRDs reveal subTAD chromosomal organization. (**A**) CRD analyses were conducted separately for each of our three ChIP-seq datasets (H3K4me3 NeuN+, H3K27ac NeuN+, H3K27ac Tissue) (see also *Figure 1A,B*). The numbers next to bars indicate the proportion of total peak population integrated into CRD structures. (**B**) Venn diagrams summarizing genome-wide sequence, in Mb megabases, integrated into CRD structures, including overlap and Jaccard similarity index between different histones and cell population (NeuN^+^ or tissue). (**C**) (top) Representative 10Mb window of chromosome 4 showing PFC NeuN+ Hi-C TAD, chromosomal loop and H3K27ac landscape including CRD structure. (bottom, shaded in gray color) Higher resolution (2Mb) peak-to-CRD assignments and peak correlational structure expressed as interaction matrix. (**D**) Genome-scale peak-to-peak correlations constrained by Hi-C loop structures. (**E**) Genome-wide base pair length distribution (x-axis log scale 10^4^-10^7^ base pairs) of CRDs (colored curves) compared to neuronal TADs (black curve). (**F**) Neuronal CTCF chromatin occupancies (Y-axis, using CTCF ENCODE reference ChIP-seq from H1 stem cell-differentiated neuronal culture) in relation to distance from CRD (colored graphs) and TAD (black graph) boundaries.

As expected, study-1 and -2 H3K27ac CRDs showed markedly higher similarity with Jaccard *J* of 0.41 as compared to cross-assay similarity between study-1 H3K4me3 CRDs and H3K27ac CRDs (*J* = 0.22) (***Figure 3B***). Furthermore, 78-79% of H3K27ac CRD peaks were putative enhancers (i.e. > 3Kb from TSS) within H3K27ac CRDs, in contrast to ∼61% within H3K4me3 CRDs (***Figure S9D***) with promoters comprising the remaining peak populations. Promoter peaks within H3K27ac CRDs were linked with an average of 3.6 enhancer peaks, compared to 1.6 enhancer peaks in H3K4me3 CRDs. Thus, the coordinated activity of histone peaks within H3K27ac CRDs comprised a higher proportion of putative enhancers, and enhancer-promoter interactions, when compared to H3K4me3 CRDs. Furthermore, as the number of regulatory elements per CRD increased, the number of enhancer-promoter interactions increases as well (***Figure S9E***). We noticed, overall, pairwise correlation between histone peaks was markedly higher within CRDs and decayed with the distance between peaks (***Figure S9F***).

Integrating neuron-specific chromosomal conformation assayed by Hi-C from NeuN^+^ sorted nuclei from PFC from a set of reference brains (N=6; 3F/3M) ^23,26^, allowed us to evaluate whether CRDs provide a submodular organization within higher-order chromatin structures such as the topologically associating domains (TADs) computed from Hi-C chromosomal conformation mappings. Examining Hi-C and H3K27ac CRD structure in the 2MB of sequence surrounding *GATB (Glutamyl-TRNA Amidotransferase Subunit B*), a locus contributing to the heritability of cognitive traits and educational attainment ^27^, reveals CRD landscapes embedded in the chromosomal TAD architecture (***Figure 3C***). We observed that pairwise correlation between histone peaks within chromosomal loop formations in the kilo- to megabase range of interspersed sequence was substantially higher than between peaks of equivalent distance but located outside of chromosomal loop contacts (***Figure 3D***). As a result, CRDs, with a median length of 120-168kb, are often sequentially organized or ‘nested’ into larger TAD domains, and frequently shared boundaries with smaller chromatin domains (‘miniTADs’) (***Figure 3E***). Furthermore, both histone CRD and TAD borders were strongly enriched for occupancies of the chromosomal scaffolding and loop organizer protein, CTCF (***Figure 3F***), which affirms that CRD modules are heavily constrained by the boundaries of their local TAD landscapes (***Figure S9G-H***).

### Acetylated CRDs show reproducible alterations in SCZ and BD PFC

Having shown that CRDs largely overlap with small, modular TADs nested within larger TADs as ‘folded-upon-self’ domains, with increased intra-TAD interactions as compared to the surrounding chromosomal portions, we examined how our case/control datasets of 739 ChIP-seq samples related to structure-function alterations of chromosomal domains in psychiatric disease at granular resolution on a genome-wide scale. 759 (13%) from a total of 6,389 H3K27ac CRDs were significantly hyper- and 587 (12%) hypo-acetylated at FDR 1% in H3K27ac NeuN+ across SCZ and control brains(***Methods, Table S7***). There was a significant correlation of the hyper- and hypoacetylated H3K27ac CRDs taken together from SCZ study-1 with the corresponding CRDs from SCZ study-2 and BD study-2 (SCZ-SCZ, ρ=0.26, P<2.2e-16: SCZ-BD ρ=0.13, P=0.051). There was also a strong correlation between SCZ study-2 and BD study-2 (ρ=0.93, P<2.2e-16) (***Figure S10***). On the other hand, none of the 2,721 H3K4me3 CRDs was significantly altered at FDR 1%, which is consistent with the peak-based analyses as described above. Overall these findings confirm that alterations in chromosomal domains are reproducible across the various postmortem studies.

### Cell type specific signatures of hyper-vs. hypoacetylated domains

To further explore the role of chromatin structure-activity in SCZ and BD, we quantified the acetylation levels of every CRD as mean of activity of histone peaks within the CRD per sample resulting in the CRD contact matrix (*m* CRDs × *n* samples) for three groups that showed disease-associated dysregulated CRDs (SCZ H3K27ac NeuN, SCZ H3K27ac Tissue and BD H3K27ac Tissue) (***Methods***). In principle, the CRD contact matrix combines CRD as a representative chromosomal structural unit (mini-TADs) with enhancer-promoter activity (***Figure 3C, Figure S9F***) from histone peaks which is in line with the underlying chromosomal HiC contact matrix but at different resolution. Indeed, principal component analysis of the CRD contact matrix (***Figure S11***) revealed strong stratification by HiC-defined A and B compartments along with hyper- and hypoacetylated CRDs across component 1 and component 2, indicating the presence of a structure-activity relationship in disease-associated CRDs. Therefore, we embarked on a more detailed analysis to explore this relationship specifically for disease-associated CRDs. Our goal was to explore an even higher order of chromatin organization by estimating CRD-CRD correlations (CRD interaction map) for the entire set of disease-associated CRDs for the SCZ groups (study-1 H3K27ac NeuN+, study-2 H3K27ac Tissue) and BD group (study-2 H3K27ac Tissue). Interestingly, the resulting CRD interaction maps were, consistently across these three groups, optimally resolved by three large clusters using K-means (***Figure 4A, Figure S12***).

**Figure 4:**
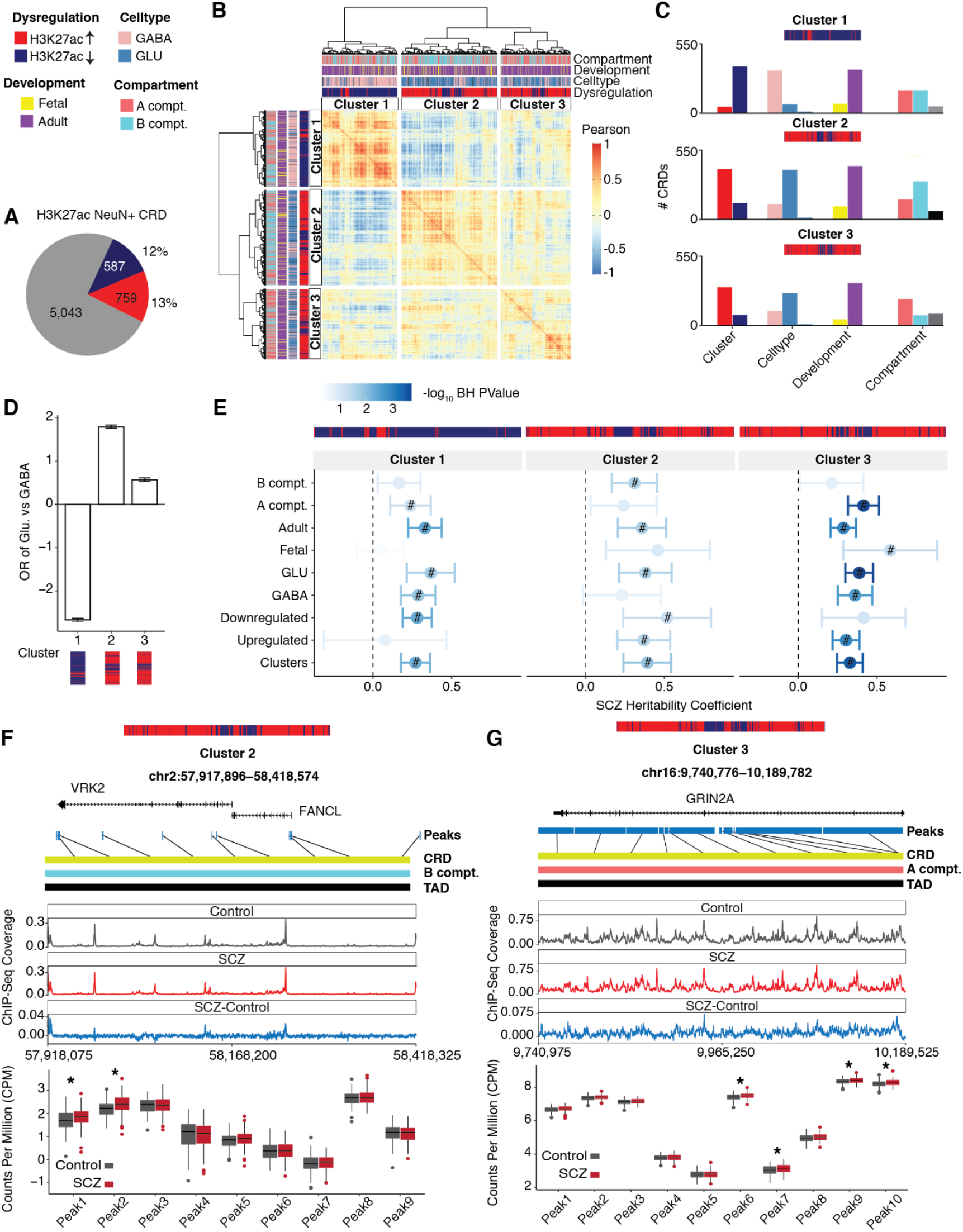
Fingerprinting disease-sensitive CRDs. Classifiers for annotated CRDs are dysregulation (hypo vs hyper-acetylation), cell type (GABA, GLU), development (fetal, adult), and compartments (A compt., B compt.) **(A)** Proportional representation of SCZ-sensitive H3K27ac NeuN+ CRDs stratified by dysregulation hypoacetylation (blue), hyperacetylation (red) and not dysregulated (gray) (**B**) K-means clustering of CRD interaction map of SCZ-sensitive H3K27ac NeuN+ CRDs into three large clusters; notice striking separation of cluster 1 representing hypoacetylated H3K27ac CRDs enriched for GABA neuron-specific peaks, and clusters 2 and 3 overwhelmingly defined by hyperacetylated H3K27ac CRDs with GLU excitatory neuron-specific peak. Cluster-specific (**C**) frequency of CRDs in every cluster stratified by every classifier (**D**) Odds ratio of enrichment of clusters in H3K27ac Gabaergic vs glutamatergic peaks using Fishers ‘s exact test. (**E**) SCZ heritability coefficients common genetic variants in SCZ by cluster and classifier. The overlap of peaks within the dysregulated CRDs in clusters with genetic variants was assessed using LD score regression. “#”: Significant for enrichment in LD score regression after FDR correction of multiple testing across all tests in the plot (Benjamini & Hochberg); “·”: Nominally significant for enrichment. Notice the overall stronger signal in cluster 3, in particular for SCZ-sensitive H3K27ac CRDs representing peaks highly acetylated in fetal CRDs. (**F**) Representative example of cluster 2 hyperacetylated CRD at *VRK2* serine/threonine kinase SCZ GWAS risk locus including (bottom) peak level changes as indicated by case (red) and control group (gray) box plots. (**G**) Representative example of cluster 3 hyperacetylated CRD at *GRIN2A* NMDA glutamate receptor neurodevelopmental risk locus (see also F).

We wondered whether such type of clustering in disease-sensitive CRD interaction map reflects a common signature in terms of spatial genome organization based on Hi-C chromosomal interaction or cell type- and developmental stage-specific regulation, or a combination thereof. To explore this, we first systematically created a resource of annotations of CRDs to a) cell types using PFC reference sample-based H3K27ac ChIP-seq resources for glutamatergic projection neurons, gabaergic interneurons and oligodendrocytes (***Table S8***) ^28^ b) developmental stages as fetal vs adult using the data for epigenetic trajectories of human cortical development ^29^, and c) chromosomal A and B compartments by utilizing the PFC NeuN+ Hi-C datasets ^23^ (***Methods, Figure S13***). Remarkably, one cluster in the CRD interaction map in three groups labeled as ‘cluster 3’(SCZ H3K27ac NeuN+ in ***Figure 4C***, SCZ H3K27ac Tissue in ***Figure S14A*** and BD H3K27ac Tissue in ***Figure S14B***), was overwhelmingly comprised of 78%-99% hyperacetylated domains harboring more glutamatergic (GLU) specific CRDs than GABAergic (GABA) CRDs. Also, cluster-3 CRDs showed much stronger enrichment for H3K27ac GLU specific peaks as opposed to H3K27ac GABA specific peaks as shown in ***Figure 4D***.

Conversely, cluster 1 of each disease group was overwhelmingly comprised (96-98%) of hypoacetylated H3K27ac CRDs, wherein >8-fold excess of GABAergic specific annotated CRDs was observed in SCZ H3K27ac NeuN+ (***Figure 4C)***. Furthermore, in H3K27ac Tissue, cluster 1 carried a large majority of oligodendrocyte-specific CRDs for both SCZ and BD (***Figure S14A-B, Figure S15***). We conclude that dysregulated acetylation of miniTAD-level chromosomal domains in PFC of SCZ and BD subjects show a striking divergence by cell type, with domain hyperacetylation consistently linked to regulatory sequences specifically important for GLU projection neurons, and hypoacetylation to inhibitory interneurons and oligodendrocytes and myelination.

### Hyperacetylated domains are linked to neurodevelopment and active chromatin status

Having uncovered strong, cell-type specific signatures among the dysregulated CRDs in our diseased PFC, we next asked whether differences in chromatin status could reveal a similar degree of stratification in the clusters of the CRD interaction map, including hypoacetylated CRDs (cluster 1), hyperacetylated CRDs (cluster 3), and mixed (hypo-/hyperacetylated) (cluster 2). Indeed, visual inspection of the CRD interaction map and CRD counts indicated A-compartment status for the majority of cluster 3 CRDs while cluster 1 and cluster 2 CRDs primarily resided in B-compartment or more mixed chromatin types (***Figure 4B-C, Figure S14A-C***). Consistent with these compartment-specific signatures, cluster 3 showed an overall higher magnitude of histone H3K27ac peak and gene expression, compared to clusters 1 and 2 (***Figure S16A-B***).

To further corroborate these cluster-specific differences in compartment types, we next explored, for each cluster in each of the three disease groups, the association of A/B chromatin organization with putative regulatory elements. To do so, we measured the enrichment of every cluster in 15 chromatin states annotated by the chromHMM Hidden Markov Model, built from adult and fetal brain-specific reference sets from REP ^30^. Interestingly, cluster 3, which was composed mainly of hyperacetylated CRDs, was selectively enriched for active chromatin types both from fetal and adult brains across the three disease groups (***Figure S17***), suggesting a link between cluster 3 CRDs and neurodevelopmental mechanisms associated with SCZ risk.

Next, we asked whether fetal CRDs (defined as CRDs in adult PFC enriched for H3K27ac fetal specific peaks ^29^, see methods for details) and A-compartment annotated CRDs show stronger enrichment of genetic variants associated with SCZ and BD in cluster 3 than clusters 1 and 2. Indeed, for each of our SCZ groups (study-1, study-2), highest enrichment by LD score regression was observed for CRDs defined by highest acetylation levels in fetal PFC. This effect was disease specific, with no enrichment found for fetal CRDs in the group of CRDs altered in BD (***Figure 4E, Figure S18A-B***), consistent with the notion that neurodevelopmental defects could be less prominent in that disorder, compared to SCZ ^31^. However, A-compartment specific CRDs in cluster-3 showed, both in the SCZ (study-1 and study-2) groups and in BD (study-2), significant LD score regression (***Figure S18B***). In addition, we noted moderate enrichment of SCZ common variants in cluster 2 comprised of both hyp- and hyperacetylated CRDs. ***Figure 4F*** illustrates, as an example, the hyperacetylated CRD encompassing the *VRK2* locus, encoding a SCZ candidate gene ^32^ important for nuclear architecture and regulation of stress-induced apoptosis in neurons ^33^.

***Figure 4G*** *shows, as* a representative example for hyperacetylated, A-compartment CRDs in cluster 3, the *GRIN2A* NMDA receptor gene locus, which carries common variants linked to SCZ risk and rare variants with high penetrance for neurodevelopmental disease ^34^. These results from our CRD-focused analyses are in concordance with our histone peak-based finding that hyperacetylation is linked to genetic variants in SCZ and BD ***(Figure S19A-B***).

### Nuclear topography of hyperacetylated domains is linked to neuronal function

Taken together, our findings show that histone CRDs are mini-TAD level-type of chromosomal domains. Therefore, we next assessed whether these clusters show differential spatial organization in the 3D genome. To examine this, we constructed a 3D genome model for adult PFC NeuN+ utilizing the TAD coordinates of PFC NeuN+ Hi-C reference sets using *chrom3D*, a Monte Carlo-type algorithm for spherical genome modeling ^35,36^. Strikingly, pairwise Euclidean distances between TAD coordinates of PFC NeuN+ that overlapped with genomic coordinates of dysregulated CRDs revealed overall significantly higher proximity and connectivity of TADs harboring cluster 3 compared to clusters 1 and 2, a 3D genome phenotype that was remarkably consistent across all three disease groups (***Methods, Figure 5A, Figure S20***).

**Figure 5:**
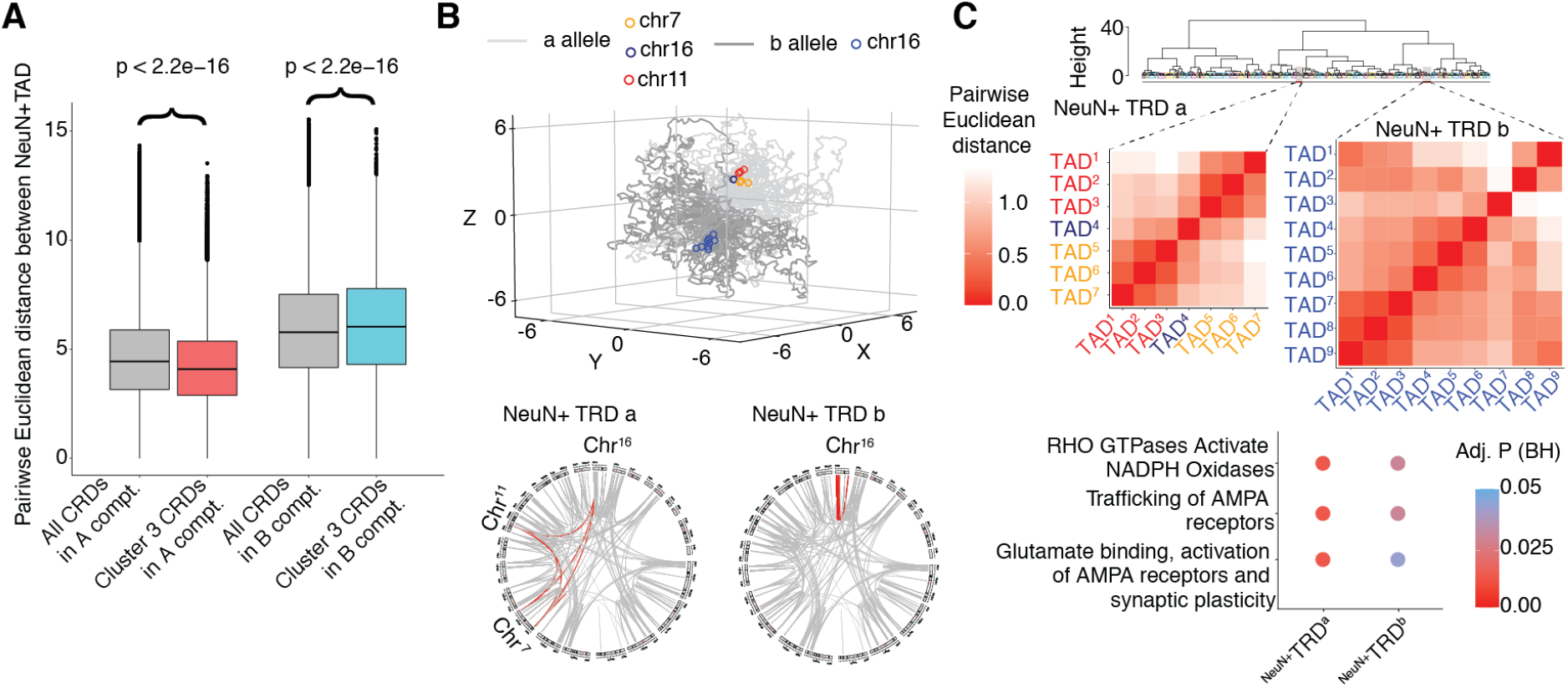
Spatial organization of cluster 3 domains in the virtual chrom3D model of the neuronal nucleus. (**A**) Box plots show pairwise Euclidean distance of A and B compartment cluster 3 H3K27ac NeuN+ CRDs, compared to genome-wide A and B compartment CRDs. (**B**) (top) chrom3D based model for spatial organization of TADs inside the neuronal nucleus. Selected TADs from chromosome 7, 11 and 16 with CRD hyperacetylation in SCZ are marked by color and share space inside the neuronal nucleus. (bottom) genome-wide circosplot for trans-chromosomal (TRD) or ultralong range (>10Mb) cis-chromosomal interactions for cluster 3 A-compartment disease-sensitive CRDs. **(C)** (top to bottom) Euclidean distance tree for cluster 3 A compartment TADs. Interaction matrix of chromosomes 7, 11, 16 from TRD shown in (B) and GO pathway enrichment for the respective CRDs.

Because clusters of disease-sensitive CRDs consistently tracked the A/B compartment-type of chromatin organization, with cluster 3 in particular overwhelmingly composed of A compartment chromatin (***Figure 5B-C, Figure S16A-B)***, we compared pairwise Euclidean distances of PFC NeuN+ TADs that overlapped specifically with A-(but not B)-compartment annotated CRDs in cluster 3. Indeed, CRDs in cluster 3 A-compartment chromatin showed significantly increased inter-TAD connectivity as observed by decreased median value of distribution of pairwise Euclidean distances compared to the distribution of pairwise Euclidean distances of genome wide annotated A-compartment CRDs (***Figure 5A-B, Figure S20***). This includes at least 80 ‘Trans Regulatory Domains(TRD)’, each comprising multiple cluster 3 TADs occupying a virtual nuclear territory of <5% spherical distance (***Methods***). Of note, 15/80 TRDs showed functional pathway enrichment, including multiple TRDs related to ion channel functions, synaptic or intracellular signaling and metabolism (***Table S9***). ***Figure 5C*** shows the diploid virtual 3D nucleome highlighting TRD no. 10 and 39, composed of multiple TADs from chromosomes 7, 11 and 16, each harboring cluster 3 domains hyperacetylated in diseased PFC. This TRD shows specific enrichment of genes regulating AMPA glutamate receptor activation and trafficking, in conjunction with RHO GTPases and NADPH oxidases and other signaling cascades linked to synaptic plasticity. These findings imply that chromosomal domain organization in the neuronal nucleus of the adult PFC goes beyond A/B compartmentalization. This includes a finer-grained 3D nucleomic architecture that results in increased chromosomal interactions, or increased virtual physical proximity in the *chrom3D* model, of domains that are at risk for hyperacetylation in SCZ and BD PFC and that share a functional signature such as the ones shown in Figure 5C, which encode genes important for glutamate receptor function.

## Discussion

The present study mapped active promoter- and enhancer-associated histone methylation and acetylation profiles in PFC of 568 brain donors, providing to date the largest histone modification dataset for SCZ and BD. Our histone peaks based analyses linked dysregulated histone acetylation predominantly to regulatory sequences for neuronal signaling and neurodevelopment, with enhancers disproportionally enriched for genetic variants carrying heritable risk for psychiatric disease. These findings strongly suggest that the epigenomic alterations in SCZ and BD brain are tracking the underlying genetic risk architecture. This outcome, when taken together with our observation that H3K27ac peak changes in rat frontal cortex exposed to long-term antipsychotic treatment showed, on a genome-wide scale, explained only (0.0005-0.0056) proportion of variance in differential SCZ and BD sensitive peaks (***Figure 1D, Figure S5G-I***), effectively rules out antipsychotic medication effects as a driver for the observed acetylation changes in our cases.

Importantly, the frontal lobe of SCZ and BD subjects is reportedly affected by alterations in histone deacetylase enzyme (HDAC) activity according to *in vivo* imaging ^37,38^, and postmortem expression studies ^39^. Likewise, transgene-derived HDAC expression in PFC neurons is detrimental for cognition and behavior ^40,41^, and negative interference with PFC HDAC expression and activity may exert therapeutic effects in animal models of psychosis 42–44.

In the second part of this study, we were able to build chromosomal domains (mini-TADs) as CRDs and CRD interaction maps by estimating the inter-individual correlations between histone peaks, which manifests higher order chromatin organization (A/B compartments). Our *in-silico* biological validation of CRDs from both assays H3K4me3 and H3K27ac clearly showed spatial clustering of histone peaks in brain epigenome. These CRDs are tightly linked to the structures of self-interacting domains, with enrichments of CTCF structural proteins at CRD domain boundaries, in line with CTCF enrichment of PFC NeuN+ Hi-C defined TAD boundaries. This particular finding could open new avenues in the genomic exploration of the brain. This is because, despite the increasing number of Hi-C reference maps from the human brain, the modular organization of chromosomes below the level of the topologically associating domains (TADs) remains poorly explored and moreover, practical considerations prohibit population level clinical studies by genome-scale Hi-C conformation mapping. In contrast, the CRD concept presented here allows for a fine-grained assessment of epigenomic alterations at several-fold higher chromosomal resolution as compared to conventional Hi-C approaches.

Our findings from clustering of disease-sensitive H3K27ac CRDs (CRD interaction map) revealed strong, cluster-specific convergences defined by A/B compartment chromatin organization, cell types and specificity towards neurodevelopment. These cluster-specific ‘fingerprints’ were consistently encountered in each of our three disease groups (SCZ study-1, SCZ study-2, BD study-2). We showed that hyperacetylated CRDs were strongly enriched for excitatory projection neuron-specific peaks with regulatory sequences indexed by chromHMM as active chromatin showing significant associations with SCZ and BD genetic risk architectures for the adult, and in case of SCZ, also the fetal PFC. Therefore, this cluster-specific fingerprint could signal that many chromatin domains important for PFC projection neurons play a critical role early in the disease process. Taken together with the notion that enhancers and other cis-regulatory sequences of the fetal brain are disproportionally over-represented among the set of common risk variants linked to schizophrenia ^3,45,46^, a finding that was replicated for the group of hyperacetylated enhancers in SCZ groups (***Figure 4E, Figure S19A***) it is plausible to hypothesize that a subset of hyperacetylated chromatin domains in diseased PFC neurons are vestiges of an early occurring neurodevelopmental disease process. Such type of epigenetic pathology in developing PFC could extend beyond the level of histone acetylation, given that alterations in DNA cytosine methylation profiles in adult SCZ PFC frequently encompass regulatory sequences defined by dynamic methylation drifts during the transition from the pre- to the postnatal period ^45^. Furthermore, the link between PFC glutamate neuron-specific CRD hyperacetylation in SCZ and BD as reported here broadly resonates with transcriptomics-focused studies reporting that in single nucleus-level expression profiling, glutamatergic neuron modules are up-regulated in the upper cortical layers of SCZ PFC ^5,47^ in addition to increased composite measures for glutamatergic transcripts in PFC of SCZ subjects ^48^. Conversely, hypoacetylated CRDs were predominantly if not overwhelmingly composed of interneuron- and oligodendrocyte specific histone peaks with no enrichment for regulatory sequences conferring heritability risk. We suggest that the majority of these domain hypoacetylation events, in line with gene expression studies proposing functional downregulation of some PFC interneuron systems and myelination pathways, reflect adaptive processes and secondary responses to the underlying mechanisms of disease.

As discussed above, hyperacetylated CRDs showed strong evidence for functional convergence, as defined by (glutamatergic) neuron-specific regulation, A-compartment chromatin organization across a wide range of development and enrichment for genetic risk. Interestingly, however, hyperacetylated CRD showed, by Hi-C spatial genome modeling, also evidence for structural convergence. This includes an overall higher inter-domain connectivity score in the *chrom3D* simulated nuclear sphere, and up to 80 ‘microclusters’ or ‘TRD’ in which hyperacetylated domains from different chromosomes share the same nuclear 3D space. Intriguingly, as exemplified by the AMPA glutamate receptor trafficking and signaling TRDs (***Figure 5C***), many of the TRDs showed enrichment for functional categories specific for neuronal function and signaling, pointing to some degree of convergence of structure, or chromosomal organization inside the nuclear sphere, and function, or neuronal signaling, at the affected domains. However, our simulation of the NeuN+ nucleus, including the Euclidean hot spots composed of hyperacetylated CRDs, ultimately represent contact frequencies in large ensembles of nuclei, not actual spatial proximities within a single nucleus. While Hi-C and chromosome conformation capture, on the 10^5^-10^6^ base pair scale largely is correlated with spatial distances in single nuclei as determined by DNA FISH ^49^, this remains to be examined for the hot spots and clustered domains discussed here.

Which types of molecular mechanisms could drive the nuclear topography of disease-relevant chromosomal domains, including the structural convergence of functionally inter-related hyperacetylated domains, as reported here? Interestingly, chromosomal contacts in brain and other tissues preferentially occur between loci targeted by the same transcription factors ^50^, with converge on intra- and inter-chromosomal hubs sharing a similar regulatory architecture including specific enhancers as well as transcription and splicing factors ^51–53^.

In summary, the work presented here describes some of the first fine-mappings of chromosomal domains in large series of disease and control brains, primarily defined by the correlational structure of nucleosomal histone modifications and then integrated into the Hi-C chromosomal conformation landscape. We expect that the findings and resources presented here, which were highly reproducible across various brain collections, will provide a unique roadmap for future studies aimed at gaining a deeper understanding of this emerging link between the neuronal 3D genome, or chromosomal organization, and cell-circuit-specific neuronal dysfunction in psychosis including underlying genetic risk architectures.

## Supporting information

SuppMethods_SuppFigures

## Acknowledgements

We thank late Pamela Sklar for her numerous contributions in the early phase of this project and Prashanth Rajarajan and Sergio Espeso-Gil for helpful discussions. This work was supported in part through the computational resources and staff expertise provided by Scientific Computing at the Icahn School of Medicine at Mount Sinai. We are extremely grateful to J. Ochando, C. Bare and other personnel of the Icahn School of Medicine at Mount Sinai’s Flow Cytometry Core for providing and teaching cell sorting expertise.

## Funding

Supported by NIH U01DA048279 and R01MH106056. PsychENCODE Consortium -- Data were generated as part of the first phase of the PsychENCODE Consortium supported by: U01MH103339, U01MH103365, U01MH103392, U01MH103340, U01MH103346, R01MH105472, R01MH094714, R01MH105898, R21MH102791, R21MH105881, R21MH103877, and P50MH106934 awarded to: Schahram Akbarian (Icahn School of Medicine at Mount Sinai), Gregory Crawford (Duke), Stella Dracheva (Icahn School of Medicine at Mount Sinai), Peggy Farnham (USC), Mark Gerstein (Yale), Daniel Geschwind (UCLA), Thomas M. Hyde (LIBD), Andrew Jaffe (LIBD), James A. Knowles (USC), Chunyu Liu (UIC), Dalila Pinto (Icahn School of Medicine at Mount Sinai), Nenad Sestan (Yale), Pamela Sklar (Icahn School of Medicine at Mount Sinai), Matthew State (UCSF), Patrick Sullivan (UNC), Flora Vaccarino (Yale), Sherman Weissman (Yale), Kevin White (UChicago) and Peter Zandi (JHU). The HBCC is funded by the NIMH-IRP through project ZIC MH002903.

## Author contributions

Wet lab work including tissue processing, sorting of nuclei and ChIP-seq and Hi-C library generation: Y.J., L.B., M.K., E.Z., R.J., J.R.W., R.P., B.S.K., L.C., O.D., S.R., J.F. Data processing and coordination: Y.J., M.K.,M.A.P., J.S.J. Bioinformatics and computational genomics: KG., G.E.H., J.B., S.R., T.G., J.P.-C., P.D. Provision of brain tissue and resources: C.A.T., S.M.,, B.K.L., D.A.L., V.H. Conception of study and design: P.R., S.A., K.G., G.E.H., J.B. Writing of the paper: K.G., S.A., P.R.

## Competing interests

The authors declare no competing financial interests.

## Data and materials availability

Raw (FASTQ files) and processed data (BigWig files, peaks, and raw / normalized count matrices) has been deposited in synapse under synID syn25705564. Browsable UCSC genome browser tracks of our processed ChIP-seq data are available as a resource at: EpiDiff Phase 2.

External validation sets used in the study are: H3K27ac ChIP-seq fetal specific peaks: Spatio-temporal enrichment of H3K27ac peaks table from http://development.psychencode.org/#, RoadMap Epigenome Project (REP) H3K27ac, H3K4me3 tissue ChipSeq peaks, chromHMM states on E073 and fetal male E081 and fetal female E082 https://egg2.wustl.edu/roadmap/data/byFileType/chromhmmSegmentations/ChmmModels/coreMarks/jointModel/final/ and CTCF ChIP-seq on human neural cell (GEO GSE127577). TruSeq3-PE.fa file was downloaded from the adaptor folder under the trimmotic repository. https://github.com/timflutre/trimmomatic/blob/master/adapters/TruSeq3-PE.fa

The source data described in this manuscript are available via the PsychENCODE Knowledge Portal (https://psychencode.synapse.org/). The PsychENCODE Knowledge Portal is a platform for accessing data, analyses, and tools generated through grants funded by the National Institute of Mental Health (NIMH) PsychENCODE program. Data is available for general research use according to the following requirements for data access and data attribution: (https://psychencode.synapse.org/DataAccess).

## Additional information

Supplementary information is available for this paper at https://doi.org/10.7303/syn25710572. Reprints and permissions information is available at www.nature.com/reprints. Correspondence and requests for materials should be addressed to P.R. or S.A.

## Notes

### Competing Interest Statement

The authors have declared no competing interest.

