## Supplementary material for "Acetylated Chromatin Domains Link Chromosomal Organization to Cell- and Circuit-level Dysfunction in Schizophrenia and Bipolar Disorder": SuppMethods_SuppFigures

**This PDF file includes:**

Methods
Captions for Supplementary Figures S1-S20
Supplementary Figs. 1-20
Captions for Supplementary Tables 1-9
References (for Methods)

**Other Supplementary Materials/ Data Files for this manuscript include the following:**

Supplementary Tables 1-9 (.xlsx)

#Correspondence to

;

;

### Methods

**Brains (postmortem):** All tissue donors of the study-1 were from the Icahn School of Medicine at Mount Sinai (MSSM), University of Pennsylvania (PENN) and University of Pittsburgh (PITT) brain bank and study-2 were from the Human Brain Collection Core (HBCC) at the national institute of mental health. Demographics of the brain cohort, toxicology and neuropathology reports are summarized in **Table 1 and Table S1**. No statistical methods were used to pre-determine sample sizes.

**Brains (rats): Chronic antipsychotic treatment**—Sprague Dawley male rats (Charles River, Wilmington, MA) were treated with a first generation (haloperidol (n=3)) or second generation atypical (risperidone (n=5) and olanzapine (n=4)) antipsychotics and placebo (n=5) in drinking water continuously for 4.5 months as previously described (1, 2). Rats were then sacrificed, trunk blood collected and plasma frozen at −20°C until analysis. Drug levels were measured as previously described [1,2] .

**ChIP-Seq library preparation and sequencing:** From the total set of 739 histone ChIP-seq datasets presented here, 28% (100 control and 109 SCZ cases) had been included in a recent PsychENCODE genomics reference paper for the adult human brain [3], the remaining 530 ChIP-seq datasets had not been presented before.

Nuclei were extracted from approximately 300mg aliquots of frozen frontal (dorsolateral prefrontal and anterior cingulate gyrus) cortex tissue, immuno-tagged with Anti-NeuN-Alexa488 (Cat# MAB377X, EMD Millipore) antibody which robustly stains human cortical neuron nuclei [4,5] for subsequent fluorescence-activated nuclei sorting. Next, chromatin of sorted nuclei was digested with micrococcal nuclease and subsequently pulled down with anti-histone antibodies, followed by library preparation and sequencing. Two histone antibodies, anti-H3K4me3 (Cat# 9751BC, lot 7; Cell Signaling, Danvers, MA) and anti-H3K27ac (Cat# 39133, Lot# 01613007; Active Motif, Carlsbad, CA) were used for immunoprecipitation. Antibody specificity was tested using peptide binding assays and immunoblotting of nuclear extracts from human postmortem cortical tissue. A commercially available histone H3 peptide array (Cat# 16–667; Millipore) containing 46 peptides representing 46 different histone H3 posttranslational modifications was used as previously described [4]. All procedures were performed as described in the recent PsychENCODE methods paper, providing a detailed description of the protocol [4]. For each cell-type specific ChIP-assay, a minimum of 400,000 sorted neuronal (NeuN+) nuclei was required as starting material. For selected gene promoters ChIP-PCR was conducted to validate cell-type specific peak profiles. Furthermore, quality controls for nuclei post-FACS included visual inspection under the microscope as described [4]. Of note, due to our stringent FACS gating criteria with maximized specificity (not sensitivity), 100% of sorted nuclei in the neuronal fraction showed green fluorescence confirming NeuN+ status, while 100% of sorted nuclei in the non-neuronal fraction only showed blue DAPI stain, confirming NeuN−status. Additional ChIP-seq studies were conducted with homogenized dorsolateral prefrontal cortex as input. To this end, frozen human postmortem brain tissue (approximately 20–200mg) was homogenized in lysis buffer and the total nuclei were purified. The nuclei solution was resuspended in 300ul of douncing buffer, treated with 2uL of micrococcal nuclease (0.2U/uL) for 5 minutes at 28 degrees Celsius, followed by 30uL of 500mM of EDTA to stop the reaction. After this initial procedure for nuclei preparation and digestion, the sample was processed in the same manner as described for the FACS sorted nuclei samples.

**Randomization and blinding:** To avoid batch effects and other confounds, samples underwent repeated rounds of randomization, including (i) chromatin immunoprecipitation procedures and (ii) library preparation. Blinding was not relevant to this study, analysts were aware of data generation, processing and donor metadata.

**Adapter sequences removal:** First the raw fastq files were corrected for adaptor pair end sequences using trimming tool called *Trimmomatic (v0.36)* [6] with the following settings: *ILLUMINACLIP:TruSeq3-PE.fa:2:30:10:8:TRUE LEADING:3 TRAILING:3 SLIDINGWINDOW:4:15 MINLEN:36.*

**Alignment, filtering, quality control and consolidation of BAM files:** Trimmed fastq files from each study were aligned to Hg38 (GRCh38) human genome using the Burrows-Wheeler Aligner (*BWA-0.7.8-r455*) method with default settings[7]. The output files were exported as BAM files. For quality control steps of BAM files, we implemented ENCODE pipeline workflow, which is as follows, 1) remove unmapped reads, mates and low quality mapping reads (mapq=30), 2) remove orphan reads and reads that were mapped to different chromosomes and 3) remove PCR duplicates using *picard* (*v2.2.4*) tool (http://broadinstitute.github.io/picard).

All BAM files from above step were tested for ENCODE quality control parameters for ChIP-Seq files: normalized strand coefficient (NSC>1.0) and relative strand coefficient (RSC>1) using *phantompeakqualtools (v2.0)* [8]. ***Figure S1A*** shows the frequency of NSC and RSC of samples from study-1 and study-2. We provide the NSC and RSC of each sample (***Data and materials availability***).

After filtering out the BAM files based on ChIP-Seq qc parameters, we prepared the files for the next step which is consolidation of bam files separately for each dataset. The objective was to subsample each ChIP-Seq library to a fixed number of mapped reads and consolidate the subsampled libraries into one file. To obtain fixed number mapped reads for subsampling of bam files, we took minimum of number of mapped reads from each study; H3K4me3 NeuN+=12M, H3K27ac NeuN+=22M and H3K27ac Tissue=23M. We obtained (median) ~30, ~60 & ~59 million of mapped paired-end reads (2 x 75bp) for H3K4me3 NeuN+, H3K27ac NeuN+ and H3K27ac Tissue respectively (***Figure S1A***). A similar procedure was followed to create a consolidated input-control file for NeuN+ study-1 and tissue study-2

**Mislabelling and contamination of samples check:** For samples mismatch and contamination check we used *QTLtools (v1.3) mbv*[9] (Match BAM to VCF) option. *MBV* takes as input a VCF file containing the genotype data for study-1 and study-2 samples and a mapped BAM file from the above section (**Alignment, filtering and consolidation of BAM files**). We did this step using the merged vcf file of genotypes of study-1 and study-2 separately. None of our samples were mismatched or contaminated. We provide a summary of *QTLtools (v1.3) mbv* [9] results of all samples (***Data and materials availability***).

**Peak Calling:** Narrow peak regions were called on a consolidated file of H3K4me3 histone mark dataset using *macs2 (v2.2.6)*[10] with Poisson p-value = 0.01 with --keep-dup all --nomodel --extsize = 150. Similarly, broad peak regions were called on study-1 and study-2 consolidated files of H3K27ac histone mark datasets using *macs2 (v2.2.6)*[10] with P-value cutoff = .01, --extsize = 150. We used a NeuN+ consolidated input control and a tissue consolidated control file separately as control inputs for peak calling on each study. All called peaks were filtered from blacklisted [11] region peaks for downstream analysis.

**Quantification of ChIP-Seq signal:** ChIP-Seq signal was quantified for every sample and every consensus peak obtained from the above section using *featureCounts (v1.5.0)* software [12]. The objective is to count the number of reads overlapping the genomic coordinates of peaks. This step results into a matrix of $m_{peaks}\ X\ n_{samples}$ 66,163 peaks X 230 H3K4me3 NeuN+, 124,054 peaks X 260 H3K27ac NeuN+ and 207,866 peaks X 249 H3K27ac Tissue, ***Tables S2A-C***

From these matrices, peaks with the low expression were filtered out using cpm of histone peaks >1 in at least 10% of samples as a threshold resulting into 64,254 peaks X 230 H3K4me3 NeuN+, 114,136 peaks X 260 H3K27ac NeuN+ and 143,092 peaks X 249 H3K27ac Tissue, ***Tables S3***. Next, the read counts were corrected for library size using the trimmed mean of M-values (TMM) method from *edgeR* library[13] and converted into the voom-normalized matrices.

**Estimation of proportion of neurons in H3K27ac Tissue:** To account for cell type heterogeneity in H3K27ac Tissue samples, we estimate the proportion of neurons using *dtangle (v2.0.9)* software[14]. Each tissue was modeled as a mixture of neurons and non-neurons. The reference samples and peaks of neurons and non-neurons were created using our previously published H3K27ac dataset on neurons and non-neurons from the PFC brain region. We provide a vector of % of neurons for each sample in the metadata table (***Data and materials availability***).

**Covariates model selection:** To estimate the technical and biological noise at sample level, we employ a 2-step approach.

1) We first identify the number of principal components using principal component analysis (PCA) method on the normalized read counts to identify the number of components that had variance of at least 1% of variance in the data. For each dataset, we take the correlation of all technical and biological covariates with the identified principal components and shortlisted the ones with FDR<20%

2) BIC: To identify the optimal number of covariates to have a good average model of histone peaks expression, we apply a Bayesian information criterion (BIC) approach[15] which introduces a penalty term for the number of parameters in the model. We start with "Diagnosis+Sex" as a base model and test all covariates one by one identified in the PCA

step. Selection criterion of a covariate in the model is at least 5% of peaks should have ($BIC_{Diagnosis+Gender+Covariate}$- $BIC_{Diagnosis+Gender}$) per histone peak >=2

Other covariates are added sequentially in this model until they fail to meet the criterion of BIC threshold. Following are the covariates that were used to correct the voom-normalized matrices for each study.

H3K4me3 $NeuN^+$ = *Sex+Diagnosis+GC content + FRIP+ Magnetic Beads*

H3K27ac $NeuN^+$ = *Sex+Diagnosis+GC content + FRIP+ Age*

H3K27ac Tissue = *Sex+Diagnosis+Neurons proportion + GC content + FRIP+*$FRIP^2$ *+ Age*

FRIP: Fraction of Reads In Peaks

***Figure S1B*** shows the distribution of variance explained by each covariate in the ChIP-Seq peaks activity matrix of each study. For a complete list of covariates see metadata table in ***Data and materials availability*** section.

**Annotating ChIP-seq histone peaks regions:** Genes and genomic Context: The Ensembl 95 genes were used for all analyses in this paper. To annotate the genomic region of a histone peak as TSS, exon, 5'UTR, 3' UTR, intronic or intergenic, we used *ChIPSeeker* (v.1.18.0) [16]. The transcript database used for the annotation is "TxDb.Hsapiens.UCSC.hg38.knownGene". We used a threshold of +/- 3kb distance from TSS of a gene for promoter annotation. ***Figure S1C*** shows the distribution of peaks annotated to categories 1) promoters, 2) introns 3) distal intergenic and 4) exon and UTRs using the hg38 transcript database imported using *ChIPSeeker* package.

**Overlap with previously published datasets:** We calculated the Jaccard index to measure the concordance of histone peaks in study-1 and study-2 with existing datasets of REP [17] and EpiMap[18]. Jaccard index is measured as the intersection of base pairs divided by union of base pairs. ***Figure S1D*** shows the pairwise similarity of datasets REP, EpiMap and study-1,2.

#### Peaks analysis

**Differential analysis:** To identify SCZ and BD sensitive peaks, we performed differential analysis on covariates corrected (**Covariates model selection section)** matrices from H3K4me3 NeuN+, H3K4me3 NeuN+ and H3K4me3 Tissue using *limma (v4.1)*[19] pipeline. ***Table S3*** provides differential analysis results from the above mentioned studies.

**Meta analysis of H3K27ac NeuN+ and H3K27ac Tissue:** Next, we combined the differential analysis results from H3K27ac NeuN+ and H3K27ac Tissue to obtain the consensus peaksets using fixed effect analysis [20]. We first created the consensus peakset by taking the set of histone peaks of H3K27ac Tissue that had at least 90% overlap (overlapping ratio = 0.90) with H3K27ac NeuN+ peaks. Then, we take the differential analysis table of overlapping peaksets of both NeuN+ and tissue to run fixed effect analysis using rma function from the R *metafor* package *(v2.0)* [20]. ***Figure S2B*** shows the results of rma analysis as a function of different overlapping ratios of H3K27ac NeuN+ and tissue peaks. The correlation plot in ***Figure S2A*** at an overlapping ratio = 0.90, shows the balance between upregulated and downregulated peaks, thus resulting in the robust fitted line passing through the origin. This becomes evident in the volcano plot of H3K27ac Meta NeuN+ shown in ***Figure S2B*** depicting the balance between upregulated and downregulated regions at an overlapping ratio =0.90.

**Pathway analysis of histone peaks:** To interpret the disease specific signatures in dysregulated H3K27ac NeuN+, Meta NeuN+ and Tissue peaks, we used the GREAT approach to assign peaks to genes. We examined the biological function of nearby genes for these non-overlapping peak regions using Genomic Regions Enrichment of Annotations Tool (GREAT) [21]. The settings for GREAT used are as follows: proximal 5.0 kb upstream, 5.0 kb downstream and plus Distal: up to 100 kb. ***Figure S4*** shows the pathway enrichment of SCZ dysregulated peaks from H3K27ac NeuN, Meta NeuN+ and Tissue and BD dysregulated peaks from H3K27ac Tissue.

**LDscore enrichment analysis:** To estimate the enrichment of brain and non-brain related GWAS in all identified histone peaks and disease sensitive peaks from H3K27ac NeuN+, Meta NeuN+ and Tissue we used *LD-score partitioned heritability (v.1.0.0)* [22]. ***Figure S6*** and ***Figure S7*** show the LDScore enrichments of SCZ and BD sensitive peaks from H3K27ac NeuN+, Meta NeuN+ and Tissue.

In LD-score partitioned heritability, it is tested if common genetic variants located in genomic regions of interest explain more of the heritability than variants not in the regions of interest, while correcting for the number of variants in either category. From this regression, a P-value as well as a regression coefficient is outputted. To enable comparisons of the regression coefficients across traits with a wide range of heritabilities, we chose to normalize it by the per-SNP heritability and named this adjusted metric the "heritability coefficient". This is not the same as the "enrichment" also outputted by the software, since the heritability coefficient takes the aforementioned baseline into account and the "enrichment" does not.

For the traits, we used the European only version of the summary statistics when available. As a consequence all GWAS results were based on individuals of European ancestry. The broad MHC-region (hg19:chr6:25-35MB) was excluded due to its extensive and complex LD structure, but, otherwise, default parameters were used for the algorithm. We ran LD-score-analyses only with sets of histone peaks covering 0.05% or more of the human genome.

**Enrichment tests of peaks in promoters and enhancers:** In this section, we test the enrichment of SCZ and BD sensitive peaks in genome wide promoters and enhancers for every study. First the genome wide peaks of H3K27ac NeuN+ and Tissue are divided into two groups 1) promoters (±3Kb from TSS) 2) enhancers (> 3Kb from TSS). The distance from TSS and promoter annotations are obtained from the previous section (**Annotating ChIP-seq histone peaks regions).** Next, we test if the fraction of SCZ or BD sensitive peaks coverage in a given study overlaps the genome wide promoters/enhancers and is significantly different from the fraction of peaks coverage by genome wide peaks as a background dataset using Fisher's Exact Test at P Value < .05. ***Figure S6A*** shows the odds ratio of enrichment tests of SCZ and BD sensitive peaks.

**Rat ChIP-Seq QC and data processing:** We followed the sample workflow as we did for human postmortem samples. For alignment of reads, we used rattus norvegicus Rnor 6.0 version of rat genome and called broad peaks using *macs2 (v2.2.6)*[10]. We did two types of differential analysis, 1) Antipsychotics vs controls: here we combine all the antipsychotic drugs into one to increase the statistical power of differential test and 2) Drugs vs controls i.e. Olanzapine vs. controls, Risperidone vs. controls and Haloperidol vs. controls separately. ***Figure S5*** shows the QC metrics from alignment and data preprocessing steps and differentials analysis results.

#### Cis-regulatory domains (CRD)

**Genome wide CRD calling:** We identify cis-regulatory domains (CRD) separately on H3K4me3 NeuN+, H3K27ac NeuN+ and H3K27ac Tissue by leveraging the inter-individual correlations of samples. Here we discuss in detail the stepwise workflow of CRD calling and identification disease specific CRDs as shown in ***Figure S8***.

**Removal of low correlation structure:** We first corrected for global effects of covariates to retain the correlation structure using PEER (probabilistic estimation of expression residuals) residualization [23] of histone peaks normalized expression from each study. A total of 18 PEER-corrected matrices $m_{peak\_PEER\_i} \times n_{samples}$ ($i = \{1, 5, 10, 15, 20, 25\}$) were produced (6 PEER-corrected H3K4me3 NeuN+, 6 PEER-corrected H3K27ac NeuN+, and 6 H3K27ac Tissue). CRDs were called on 18 matrices individually using the following R functions from the *decorate (v1.0.14)* [24] package.

$$CRDlist = runOrderedClusteringGenome(m_{peak\_PEERj} \times n_{samples}, peaks_{genomiCoordinates}, method.corr = "spearman")$$

$$CRDClusters = createClusters(CRDlist, method = "meanClusterSize", meanClusterSize = c(10, 25, 50, 80, 100))$$

$$CRDScore = scoreClusters(CRDlist, CRDClusters)$$

The output from above mentioned commands were 18 *CRDScore* objects (6 for each study datasets) containing a table of histone peaks assigned to CRDs, their mean correlation, and lead eigen factor (LEF). LEF of a CRD is a fraction of variance explained by the first eigenvalue of the correlation matrix $[m \times m]$ of histone peaks located within a CRD. Larger LEF values (i.e. >10%) can be interpreted as strongly correlated peaks whereas smaller values correspond to weaker correlations of peaks located within a CRD. Filtering out the CRDs with weaker correlations is an important step because it substantially reduces the burden of multiple testing in differential CRD analysis.

**CRD filtering and merging:** To filter out CRDs with weaker correlations, histone peaks positions were shuffled per chromosome for all samples to create permuted matrices $m_{Permutation\_j\_peaks\_PEER\_i} \times n_{samples}$ (where $i = \{1, 5, 10, 15, 20, 25\}$ and $j=1\text{-}10$). A total of 180 PEER-corrected matrices $m_{peak\_PEER\_i} \times n_{samples}$ ($i = \{1, 5, 10, 15, 20, 25\}$) were produced (10 permutations × 6 PEER-corrected matrices × 3 datasets; $m_{Permutation\_j\_peaks\_PEER\_i} \times n_{samples}$) .

CRDs were called on 180 matrices individually using the following R functions from the *decorate* package.

CRDs calling on 180 matrices (10 permutations × 6 PEER-corrected matrices × 3 datasets; $m_{Permutation\_j\_peaks\_PEER\_i} \times n_{samples}$) on permuted matrices followed the same workflow as explained. Lastly, $LEF_{cutoff}$ was obtained as vectors were combined from $10 CRDScore$ lists obtained from CRD calling on $m_{Permutation\_j\_OCR\_PEER\_i} \times n_{samples}$ ($i$ =*1-10*)

$$LEF_{Permuted} = \left[LEF_1, LEF_2, LEF_3, ..., LEF_{10}\right], where\ LEF_i = N \times 1$$

$$LEF_{cutoff} = Pr(LEF_{Permuted} = threshold),\ where\ threshold = 1 - 0.10$$

Final table of CRDs was obtained by keeping all CRDs with $LEF_{measured} > LEF_{cutoff}$ in $CRDScore$ list of $m_{peaks\_PEER\_i} \times n_{samples}$. ***Figure S9A*** shows an example of distribution of $LEF_{measured}$ and $LEF_{permuted}$.

Next, overlapping CRDs of different sizes were merged to obtain discrete CRDs for downstream analysis. To decide the optimal number of PEER factors, we measured the $LEF_{cutoff}$ of CRDs called on the input matrix histone peaks matrix residualized by various numbers of PEER factors; we tested *{1, 5, 10, 15, 20, 25}* PEER factors (***Figure S9B***). The the number of peaks within the CRDs are shown in ***Figure S9C***, while the final lists of coordinates of CRDs of study-1 H3K4me3 NeuN+, H3K27ac NeuN+ and H3K27ac Tissue are provided in **Table S6**.

**In-silico biological validation of CRDs:** To validate 3D interactions captured by CRDs with Hi-C dataset, we used CTCF ChIP-seq peak list from ENCODE human neural cells [25] (***Data and materials availability***). We quantified the density of CTCF sites in 200 bins (each bin size equals to 1kb) around CRD boundaries (***Figure 3F***). In order to quantify how many *in-silico* 3D interactions captured by CRDs are within the 3D interactions measured as Topologically Associated Domains (TADs) from PFC NeuN+ Hi-C experiments [26], we measured the number of CRDs overlapping with PFC NeuN+ Hi-C TADs stratified by number of TADs (*N ={0,1,2,3,>=4}*). Next, we measured how many PFC NeuN+ Hi-C TADs are within the CRDs stratified by the number of CRDs (*N ={0,1,2,3,>=4}*). We measured the correlation of histone peaks inside the Hi-C loops and outside the Hi-C loops to show peaks inside the Hi-C loops have more correlation than the peaks outside the Hi-C loops.

**Glutamateric, GABAergic, and oligodendrocyte ChIP-seq data:** The data were obtained from [27]. H3K27ac peaks were called with DFilter [28] using the following parameters: "-f=bam -pe -ks=60 -lpval=4". For each cell type, H3K27ac peak lists for replicate samples were then overlapped using a custom R script, and peaks which were present in at least half of replicates were preserved for further analysis (peak numbers: 44,519 in GABA neurons, 46,580 in Glu neurons, 45,963 in OLIG cells). Peaks detected in Glu, GABA and OLIG cells were further overlapped using the bedtools package to obtain Glu-specific (19,697), GABA-specific (16,297), and OLIG-specific (26,975) peaks (***Table S8***).

**Annotation of CRDs:** CRDs annotation to 1) development category as fetal and adult [29], 2) cell types as glutamatergic (GLU), gabaergic (GABA) and oligodendrocytes (OLIG) from (***Table S8***) [27,30], 3) active compartment as A and outside A compartment and 4) inactive compartment as B and outside B compartment using the PFC NeuN+ HiC data [26]. Every CRD was assigned to a specific category, if the fraction of peaks coverage in a CRD in a given assay matches the testing dataset (using the data resources as explained above) and is significantly different from the fraction of peaks coverage in all other CRDs as a background dataset using Fisher's Exact Test at Pvalue < .05. For chromosomal environment annotation (3), a fraction of full CRD coverage was used for annotation instead of using coverage of peaks within the CRD. ***Figure S13*** shows the final counts of CRDs annotated to each category of cell type, development and compartments and ***Table S7*** lists the annotation of CRDs from three disease groups.

**Disease specific CRDs:** In this section, we show how differential analysis of CRDs was done and how we identified the relationship between structure and activity of CRDs.

**Differential CRD analysis:** To perform differential CRD analysis to identify SCZ and BD sensitive CRDs, we used the peak differential analysis table as input (**Table S3**). Here, we show the calculation of $P$-value and $\log_2$(fold change) for one CRD ($CRD_x$) that is linked to $k$ Peaks.

$$CRD_x = [Peak_1, Peak_2, Peak_3, ..., Peak_k], \text{ where } CRD_x \text{ has } k \text{ histone peaks}$$

$$M = min\left\{Pvalue_{Peak_{=i}}\ ;\ i = 1, 2, 3, ..., k\right\}$$

$$Pvalue_{CRD_x} = 1 - \left(1 - M^k\right)$$

$$log_2FC_{CRD_x} = \frac{1}{k}\sum_{i=1}^{k} log_2FC\left(Peak_i\right)$$

Final differential analysis table is obtained after applying FDR correction on genome wide $Pvalue_{CRD_x}$ vector using the p.adjust R function with "fdr" option.

**CRD interaction map:** Next we quantified the expression of CRD for m $_{\text{CRDs}}$ and n $_{\text{samples}}$ as CRD contact matrix by taking the mean of peaks that are within the CRD per sample as shown in the equation below.

$$CRD\ contact\ matrix = \sum_{i=1}^{k} CPM_{peaks}$$

We applied K-means clustering [31] on correlation of disease sensitive CRD contact matrix and preselected K=3. The K=3 threshold was chosen based on uniform distribution of A-compt and B-compt across clusters CRD interaction map of SCZ sensitive H3K27ac NeuN+, H3K27ac Tissue and BD sensitive H3K27ac Tissue.

**LDscore enrichment analysis of CRDs:** To estimate the enrichment of brain and non-brain related GWAS in all identified CRD and disease sensitive CRD, we tested the genomic regions of histone peaks within the CRDs from H3K27ac NeuN+, Meta NeuN+ and Tissue and applied *LD-score partitioned heritability (v.1.0.0)* [22] as explained in LDscore enrichment analysis of peaks section. ***Figure S18*** and ***Figure S19*** show the LDScore enrichments of SCZ and BD sensitive CRDs from H3K27ac NeuN+ and H3K27ac Tissue.

**Enrichment tests of CRDs in development, cell type and chromatin organization:** We test the enrichment of peaks in CRDs in cell types (***Figure 4D, Figure S15***), development (***Figure S17***) and A and B compartment chromatin organization (***Figure S16***). Each of the enrichment tests was done as explained in **Enrichment tests of peaks in promoters and enhancers** before.

**Modeling chromatin conformation in 3D:** Hi-C data from PFC NeuN+ was used to infer chromatin conformation structure in 3D. We used bulk Hi-C data from PFC NeuN+ from four adults. Primary processing was performed with the HiC-Pro pipeline [32] at 50kb and 1Mb resolution. In order to improve the accuracy of 3D modeling, we combined data from different donors for the PFC NeuN+ to increase sequencing depths. Contact matrices produced by HiC-Pro were converted to cooler format using HiCExplorer [33], balanced using the cooler suite of tools [34], excluding the ENCODE v3 blacklisted regions (https://www.encodeproject.org/files/ENCFF356LFX/) from balancing with the `cooler balance --blacklist` parameter. Topologically associated domains (TADs) were called at 50kb using the `diamond-insulation` algorithm implemented in the cooltools suite (https://cooltools.readthedocs.io/en/latest/). Hi-C contact matrices and TAD calls were preprocessed to `gtrack` files as input to Chrom3D, as previously described [35,36]. For more details on HiC data generation and processing see methods section on HiC [26]. We restricted our analysis to diploid autosomal interactions, 50kb for intrachromosomal and 1Mb for interchromosomal. Chrom3D was run with nucleus radius of 5.0 for 2E6 iterations, `--radius 5.0 --iterations 2000000`. XYZ-coordinates were parsed from the output `cmm` files.

**Trans regulatory domain analysis**: We took the coordinates of SCZ sensitive H3K27ac NeuN+ and Tissue CRDs and BD sensitive H3K27ac Tissue CRDs and overlapped with the PFC NeuN+ TAD coordinates obtained from the **Modeling chromatin conformation in 3D** section. To test the presence of localized SCZ or BD sensitive CRDs in 3D genome, we measured the pairwise 3D distance of the TADs that overlapped with diseased CRDs stratified by clusters 1,2 and 3 as shown in ***Figure 20.*** Cluster 3 was clearly the cluster with most localized TADs than cluster 1 and 2. To further explore the localization of domains of TADs from cluster 3, we did hierarchical clustering of pairwise 3D distance of the TADs to identify the trans regulatory domains that could be associated with specific pathways. Figure 5C shows an example of two TRDs from H3K27ac NeuN+ that were replicated in H3K27ac Tissue in SCZ. We ran the pathway analysis on CRDs overlapping with the TADs that are within the TRDs.

### Supplementary Figures

**Figure S1** | Mapped reads and phantompeakqualtools quality metrics.

**Figure S2** | H3K27ac Meta NeuN+ differential analysis.

**Figure S3** | Concordance analysis between the SCZ and BP effect sizes across and within two studies.

**Figure S4** | Pathway analysis of SCZ and BD dysregulated peaks.

**Figure S5** | Quality metrics of rat H3K27ac Tissue study.

**Figure S6** | Enrichment of SCZ dysregulated peaks in risk variants of psychiatric and non-psychiatric diseases stratified by promoters and enhancers.

**Figure S7** | Enrichment of BD dysregulated peaks in risk variants of psychiatric and non-psychiatric diseases.

**Figure S8** | Schematic workflow of the steps in CRD analyses.

**Figure S9** | CRD filtering and merging.

**Figure S10** | Concordance between sets of dysregulated CRDs across and within the studies.

**Figure S11** | Demonstration of chromatin structure-activity relationship in CRDs.

**Figure S12** | CRD interaction Map.

**Figure S13** | Annotated CRDs.

**Figure S14** | Clusters of disease specific CRD interaction map.

**Figure S15** | Enrichment of disease specific H3K27ac Tissue CRDs in oligodendrocytes.

**Figure S16** | Spatial organization of disease specific CRDs.

**Figure S17** | REP ChromHMM annotations of the DLPFC adult and fetal brain genome.

**Figure S18** | Cluster 3 is strongly enriched for SCZ and BD risk variants.

**Figure S19** | Cluster 3 is strongly enriched for risk variants associated with psychiatric traits.

**Figure S20** | Three dimensional distance of diseased TADs in a three dimensional genome.

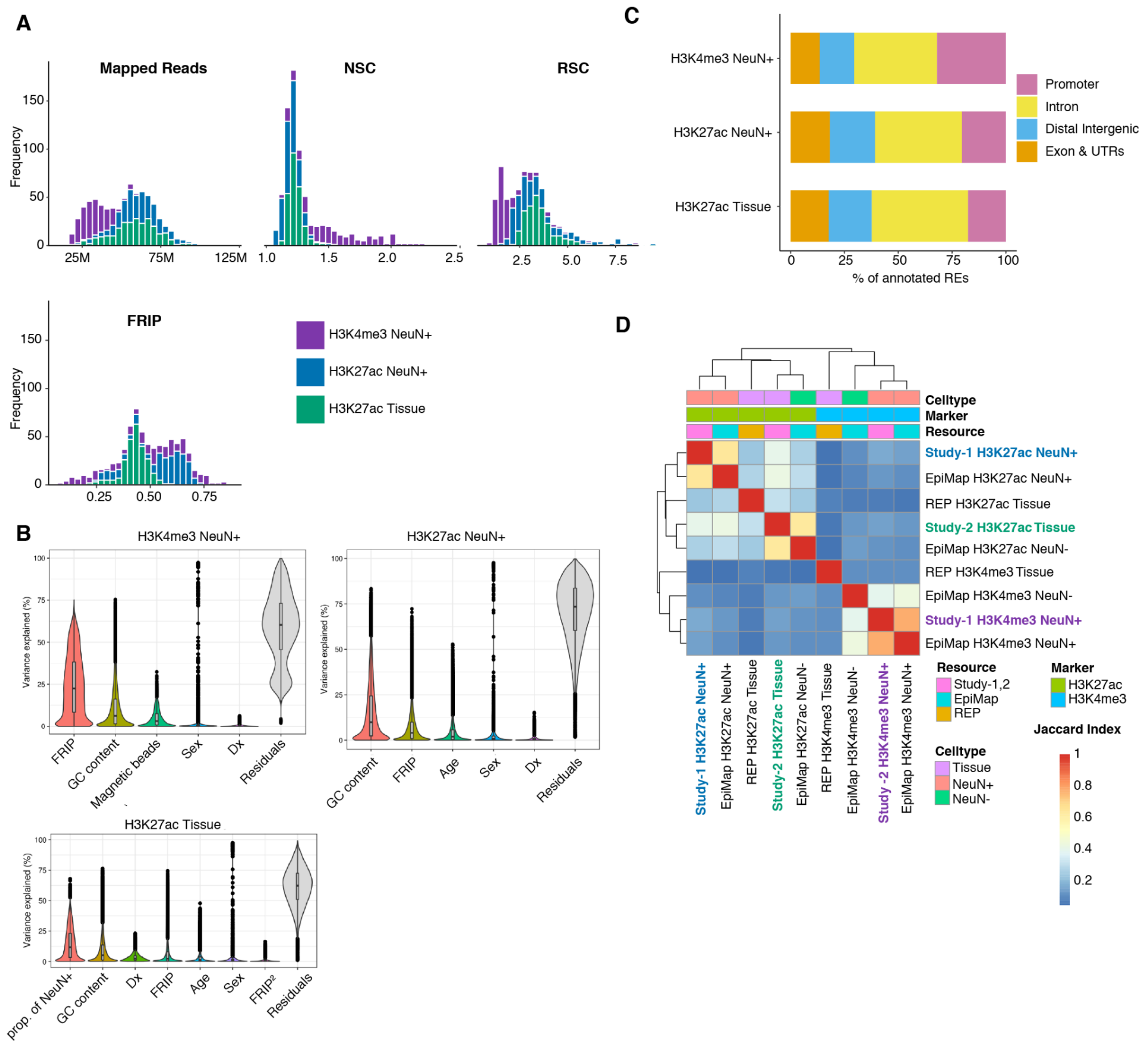


**Figure S1 | Mapped reads and phantompeakqualtools quality metrics. (A)** The distribution of mapped reads, Normalized Strand Cross-correlation coefficient (NSC) distribution, Relative Strand Cross-correlation coefficient (RSC) and Fraction of Reads In Peak (FRIP) for 230, 260 and 249 samples stratified by studies (H3K4me3 NeuN+, H3K27ac NeuN+ and H3K27ac Tissue in purple, blue and green, respectively). **(B)** Violin plots to show the distribution of variance in each peak explained by the identified confounds plotted as x-axis across all three studies. **(C)** Bar plot to show the percentage of peaks annotated to categories 1) promoters in pink, 2) introns in yellow, 3) distal intergenic in light blue and 4) exon and UTRs in orange using the grch38 transcript database imported using chipseeker package. **(D)** Heatmap to show Jaccard index between pairs of datasets 1) study-1 H3K27ac NeuN+, 2) EpiMap H3K27ac NeuN+, 3) REP H3K27ac Tissue, 4) study-2 H3K27ac Tissue, 5) EpiMap H3K27ac NeuN-, 6) study-1 H3K4me3 NeuN+ and 7) EpiMap-H3K4me3 NeuN+.

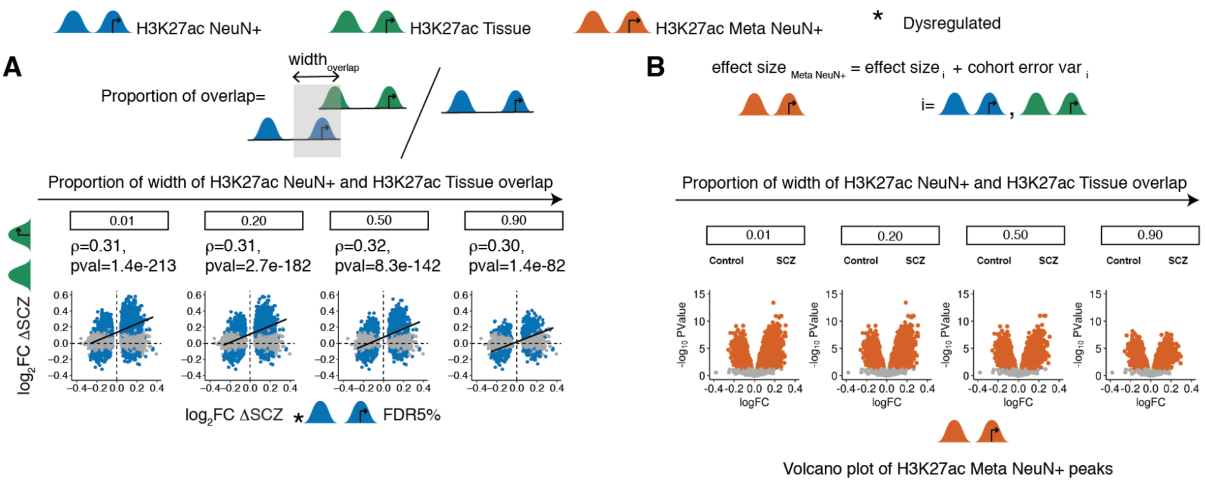


**Figure S2 | H3K27ac Meta NeuN+ differential analysis.** Top row shows the representation of H3K27ac NeuN+, H3K27ac Tissue and H3K27ac Meta NeuN+ histone peaks in blue, green and orange respectively. Asterisk sign shows dysregulated peaks obtained after differential analysis **(A)** Illustration of how proportion of overlap was measured between H3K27ac NeuN+ and H3K27ac Tissue peaks. For example, for a H3K27ac Tissue peak that overlaps with H3K27ac NeuN+ peak, the width of the overlapping region is divided by the total width of H3K27ac NeuN+ peak to get the proportion of overlap. Correlation plot of $\log_2$FC (SCZ vs. controls) H3K27ac NeuN+ at FDR 5% from study-1 with $\log_2$FC (SCZ vs. controls) H3K27ac Tissue from study-2 between the set of overlapping peaks across two studies as a function of increasing proportion of overlap. **(B)** Illustration of the effect size of H3K27ac Meta NeuN+ as summation of the effect sizes from the overlapping peaksets of H3K27ac NeuN+ and H3K27ac Tissue and an error term that accounts for differences across two studies. Volcano plot of $\log_2$FC (SCZ vs controls) H3K27ac Meta NeuN+ vs -$\log_{10}$(PValue) from fixed effect analysis of peaksets as a function of increasing proportion of overlap.

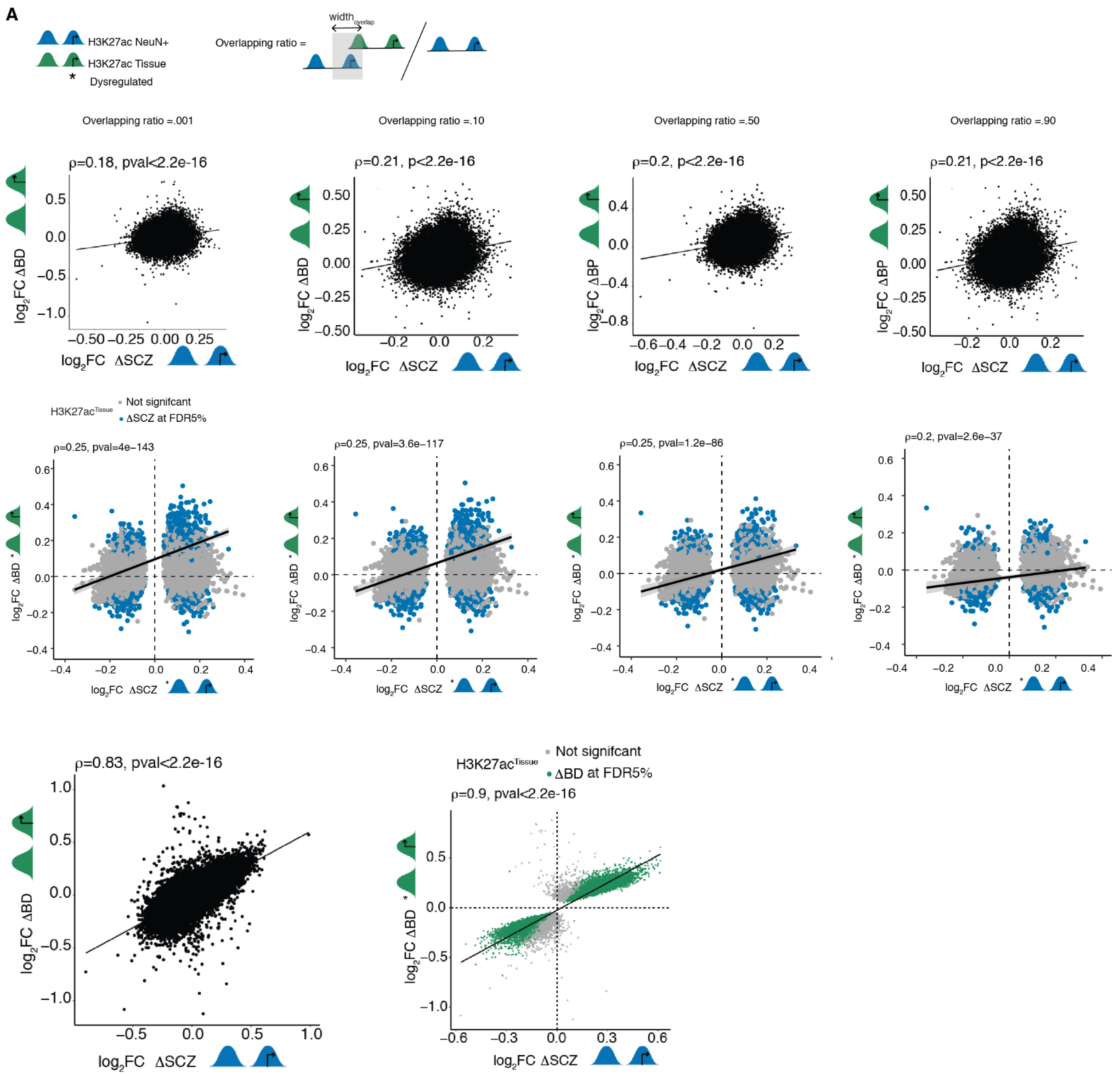


**Figure S3 | Concordance analysis between the SCZ and BP effect sizes across and within two studies.** Top row shows the representation of H3K27ac NeuN+ and H3K27ac Tissue histone peaks in blue and green respectively. Asterisk sign shows dysregulated peaks obtained after differential analysis along with an illustration of the overlapping ratio of H3K27ac NeuN+ and H3K27ac Tissue peaksets. First row shows spearman correlation between $\log_2$FC (SCZ vs. controls) H3K27ac NeuN+ and $\log_2$FC (BD vs. controls) H3K27ac Tissue and second row shows correlation between $\log_2$FC (SCZ vs. controls) H3K27ac NeuN+ at FDR 5% and $\log_2$FC (BD vs. controls) H3K27ac Tissue. The bottom row is the correlation plot within the studies i.e. $\log_2$FC (SCZ vs. controls) H3K27ac Tissue and $\log_2$FC (BD vs. controls) H3K27ac Tissue.

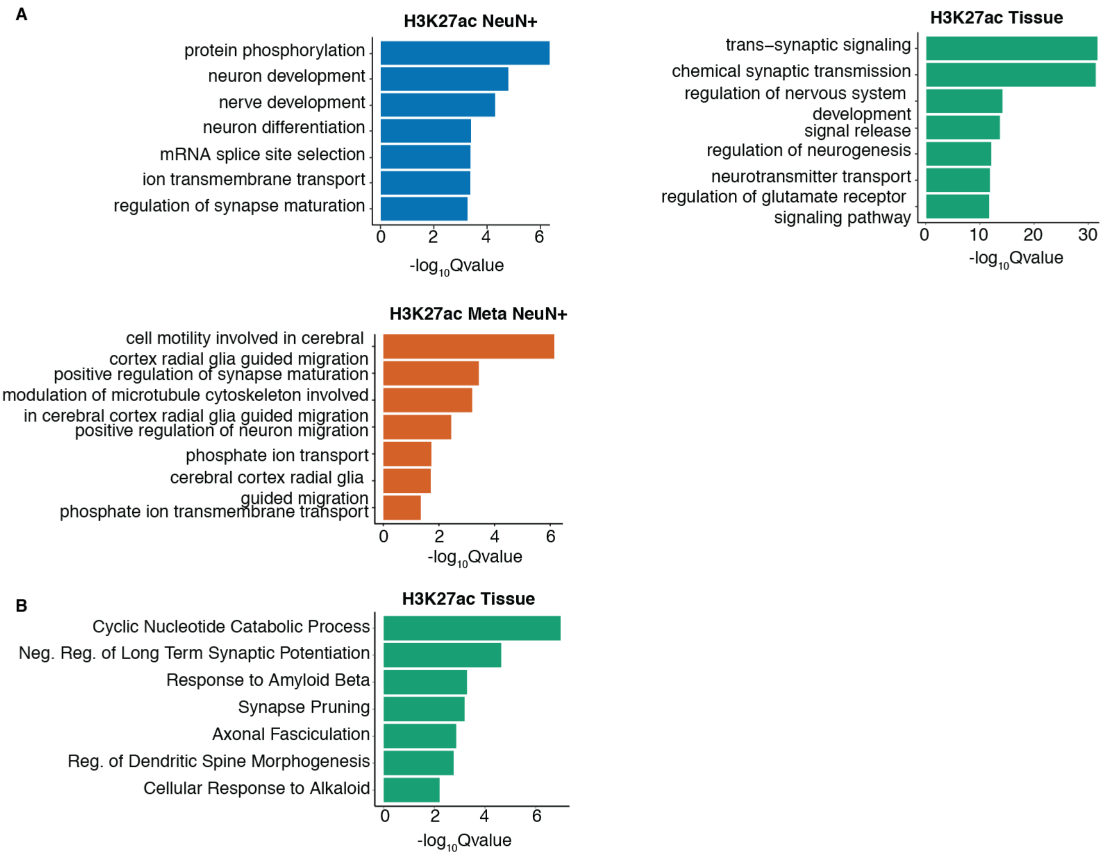


**Figure S4 | Pathway analysis of SCZ and BD dysregulated peaks. (A)** Gene set enrichment analysis of dysregulated peaks at FDR 5 % identified from differential analysis of SCZ vs controls in H3K27a NeuN+, H3K27ac Tissue and H3K27ac Meta NeuN+. **(B)** Gene set enrichment analysis of dysregulated peaks at FDR 5 % identified from differential analysis of BD vs controls in H3K27ac Tissue.

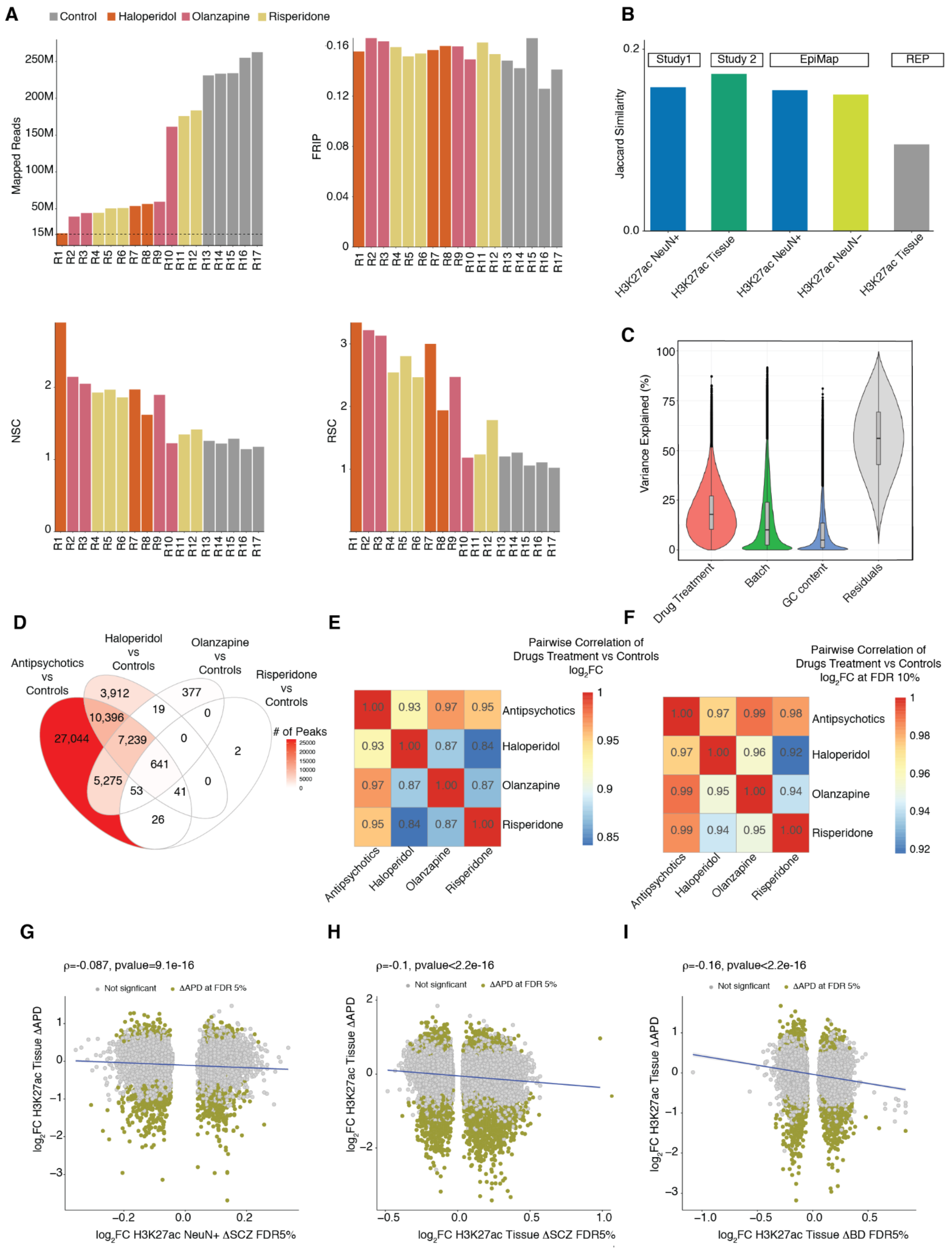


**Figure S5 | Quality metrics of rat H3K27ac Tissue study. (A)** Bar plot of mapped reads, Normalized Strand Cross-correlation coefficient (NSC) distribution, Relative Strand Cross-correlation coefficient (RSC) and Fraction of Reads In Peak (FRIP) for Rats with (n=4) olanzapine, (n=3) haloperidol, (n=5) risperidone antipsychotic drug treatments and (n=5) vehicle controls. **(B)** Bar plot of Jaccard index shows the similarity between Rat H3K27ac Tissue peaks after liftover to hg38 and study 1 H3K27a NeuN+, study 2 H3K27ac Tissue, EpiMap H3K27ac NeuN+ and REP H3K27ac Tissue. **(C)** Violin plot shows the distribution of variance in each peak explained by the identified confounds plotted as x-axis. **(D)** Venn diagram shows the overlap between the pairs of dysregulated peaks at FDR 10% after differential analysis across following categories: 1) Antipsychotic (olanzapine+haloperidol+risperidone) vs. controls, 2) olanzapine vs controls, 3) haloperidol vs controls and 4) risperidone vs. controls. **(E)** Heatmap of *pi1* statistics estimated as the proportion of non-null tests performed on a vector of differential peaks nominal p values and **(F)** only those p values that pass the cutoff of FDR 5% of each group out of four aforementioned groups in other groups. Spearman correlation between $\log_2$FC (Antipsychotics drugs vs. controls) H3K27ac Rat Tissue with **(G)** $\log_2$FC (SCZ vs. controls) H3K27ac NeuN+ , **(H)** $\log_2$FC (SCZ vs. controls) H3K27ac Tissue and **(I)** $\log_2$FC (BD vs. controls) H3K27ac Tissue at FDR 5%

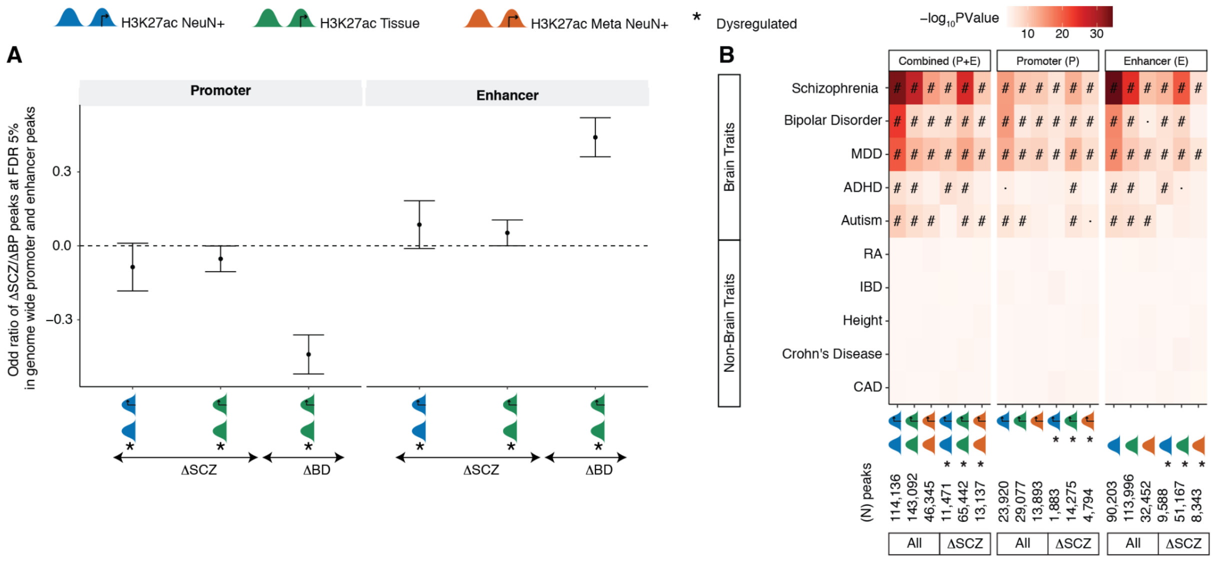


**Figure S6 | Enrichment of SCZ dysregulated peaks in risk variants of psychiatric and non-psychiatric diseases stratified by promoters and enhancers.** Top row shows the representation of H3K27ac NeuN+, H3K27ac Tissue and H3K27ac Meta NeuN+ histone peaks in blue, green and orange respectively. Asterisk sign shows dysregulated peaks obtained after differential analysis. **(A)** Odds ratio of enrichment of dysregulated peaks at FDR 5% from differential analysis (SCZ vs. controls and BP vs. controls) in genome wide promoters (left) and enhancers (right) with genome wide peaks as background of H3K27ac NeuN+ and H3K27ac Tissue. **(B)** Heatmap of enrichment *P*-values for brain and non-brain related GWAS traits overlapping with histone peaks from study-1 H3K27ac NeuN+, study-2 H3K27ac Tissue and study-1 H3K27ac Meta NeuN+. All dysregulated peaks in (B) are specific to SCZ. The heatmap in B) shows stratification on the top by 1) combined (P+E) promoters and enhancer peaks taken together, 2) promoters only (peaks within ±3Kb from TSS) and 3) enhancers (peaks other than promoters) peaks. Each of this category is further stratified into 1) all peaks and 2) dysregulated peaks at FDR 5% after differential analysis (SCZ vs. controls) across all three groups H3K27ac NeuN+, H3K27ac Tissue and H3K27ac Meta NeuN+ . "#": Significant for enrichment in LD score regression after FDR correction of multiple testing across all tests in the plot (Benjamini & Hochberg); "·": Nominally significant for enrichment.

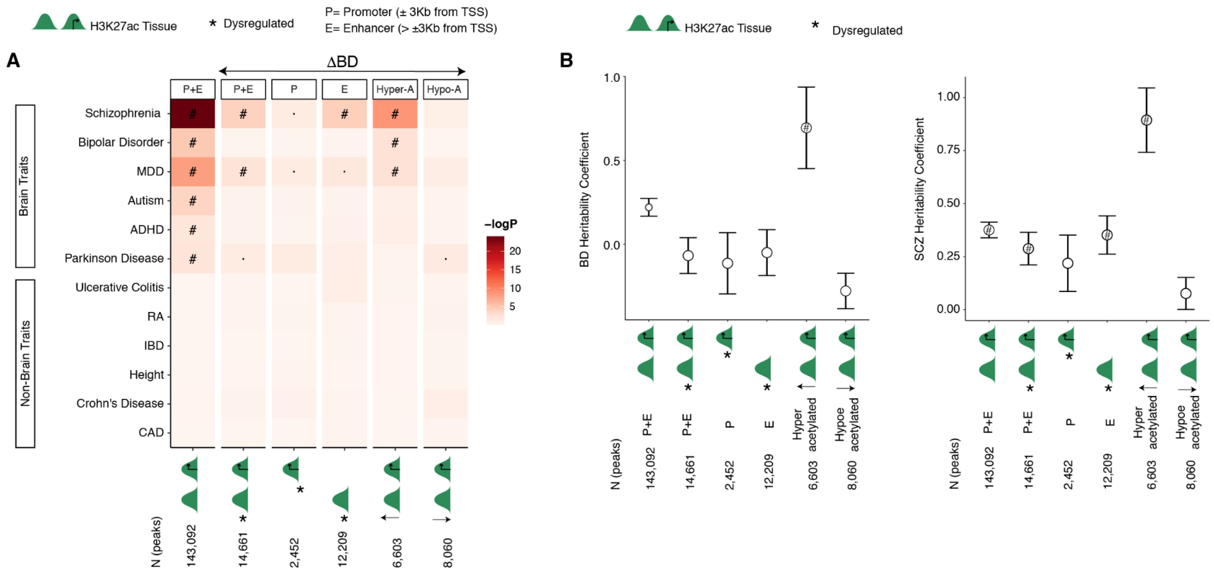


**Figure S7 | Enrichment of BD dysregulated peaks in risk variants of psychiatric and non-psychiatric diseases.** Top row shows the representation of H3K27ac Tissue histone peaks in green and an asterisk sign to show dysregulated peaks obtained after differential analysis. P and E abbreviations are explained as promoters are peaks that are within ±3Kb from TSS and enhancers are the ones that are located > ±3Kb from TSS. **(A)** Heatmap of enrichment *P*-values for brain and non-brain related GWAS traits overlapping with histone peaks within BD sensitive H3K27ac Tissue peaks. The heatmap in B) shows stratification on the top by 1) (P+E) combined promoters and enhancer peaks taken together, 2) (P+E) dysregulated peaks, 3) (P) dysregulated promoters (peaks within ±3Kb from TSS), 4) (E) dysregulated enhancers (peaks other than promoters), 5) (Hyper-A) hyperacetylated ($log_2FC>0$ and adjusted P value < 5%) and 6) (Hypo-A) hypoacetylated ($log_2FC<0$ and adjusted P value < 5%). **(B)** BD (left) and SCZ (right) heritability coefficients of common genetic variants in BD overlapping histone peaks from above mentioned 6 categories. The overlap of peaks with genetic variants was assessed using LD score regression. ”#”: Significant for enrichment in LD score regression after FDR correction of multiple testing across all tests in the plot (Benjamini & Hochberg); ”·”: Nominally significant for enrichment.

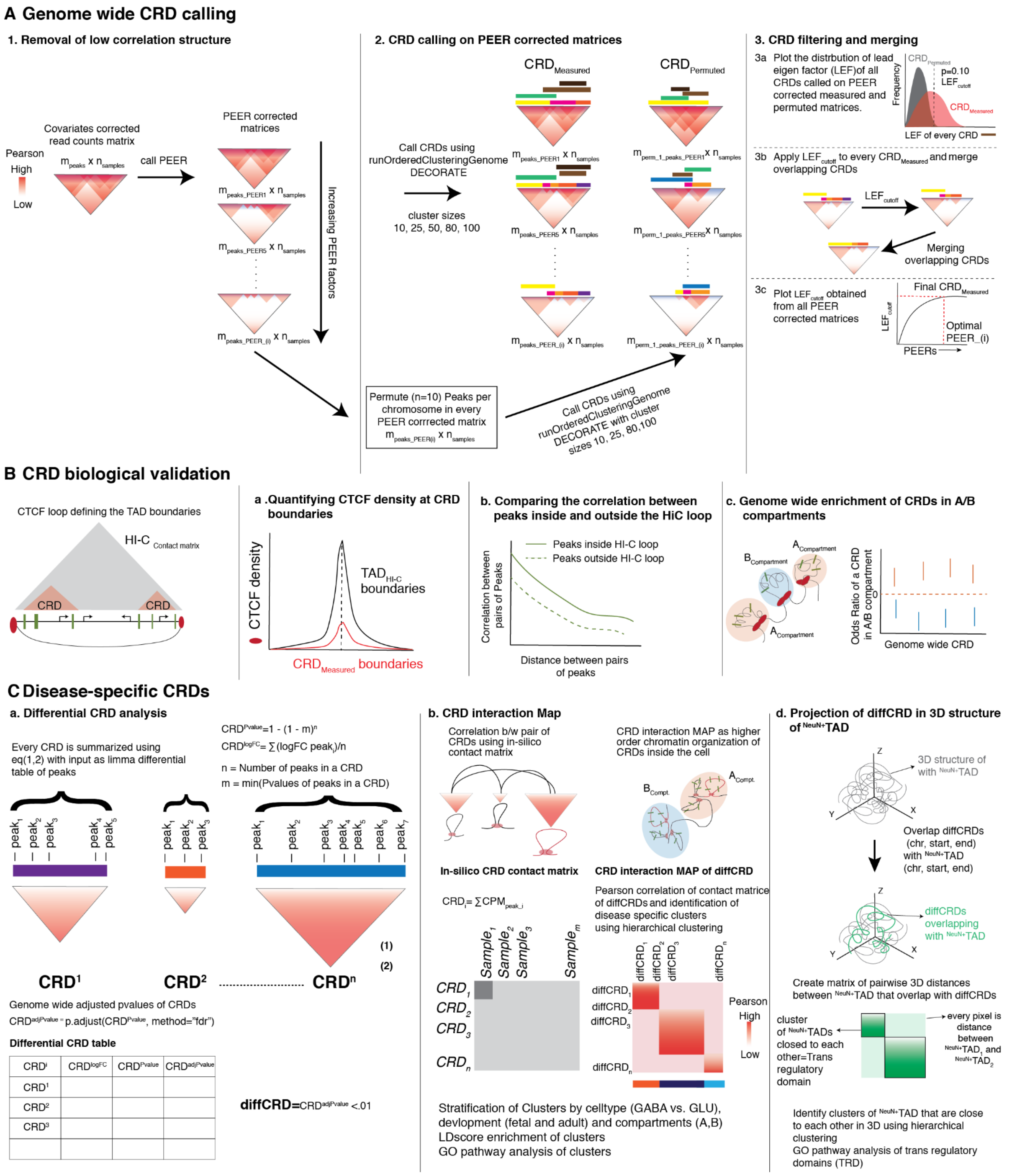

**Figure S8 | Schematic workflow of the steps in CRD analyses. (A)** Genome wide CRD calling includes data preprocessing, genome wide CRD calling and filtering and merging of CRDs. **(B)** *In-silico* biological validation of CRDs including the CTCF enrichment at the boundaries of CRDs, comparison of correlation of peaks that are inside and outside the HiC-loop and enrichment of CRDs in A and B compartments .**(C)** Identification of disease specific CRDs including differential analysis of CRDs, CRD interaction map from correlation of CRD contact matrix and finally projection of identified dysregulated CRDs in 3D genome. 3D genome was simulated using *chrom3D*.

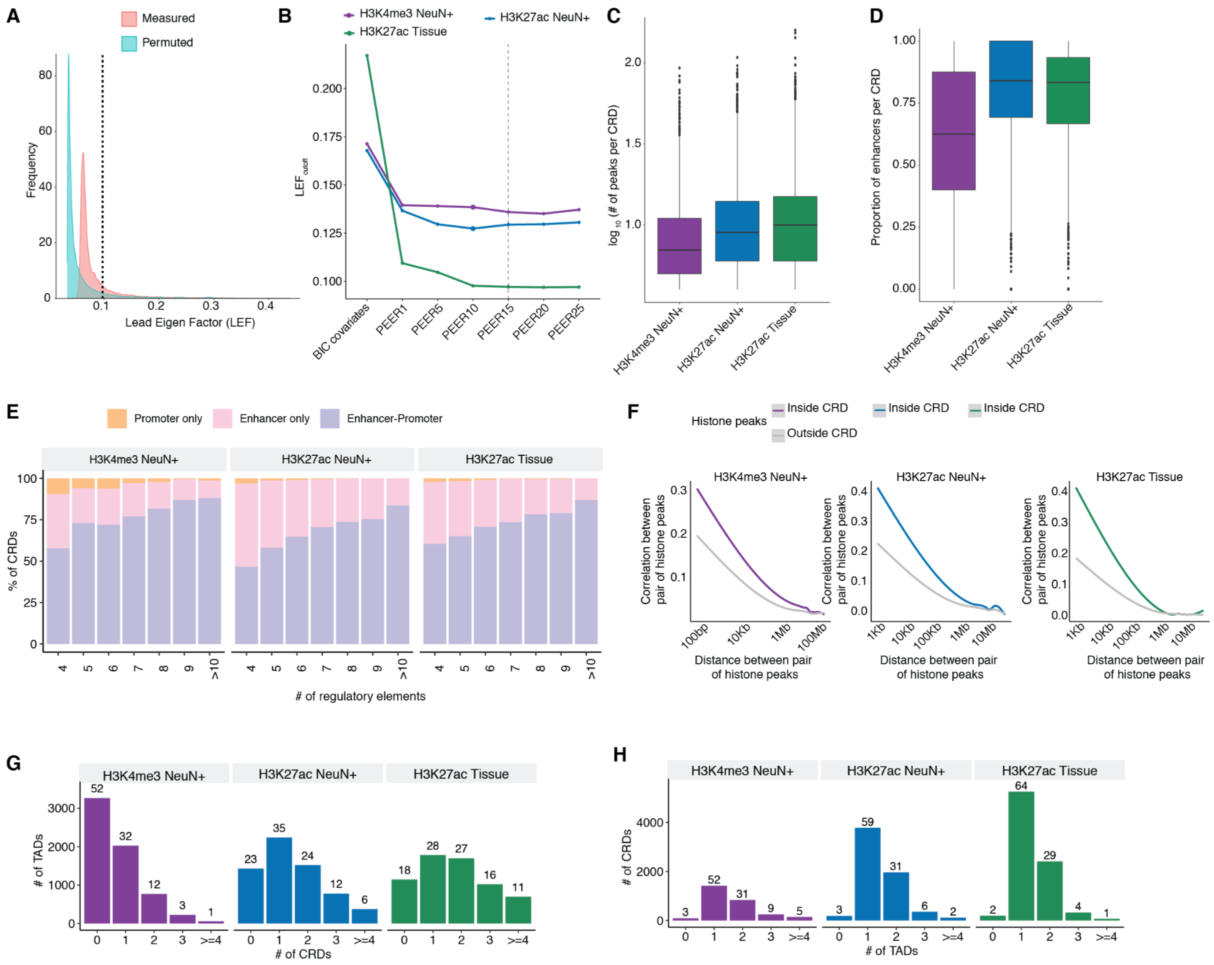


**Figure S9 | CRD filtering and merging. (A)** An example illustrating filtering step using distribution of Lead Eigen Factor (LEF) of CRDs called on $m_{peaks\_PEER} \times n_{samples}$ and $m_{Permutation\_i\_OCR\_PEER} \times n$ samples ($i=10$ permutations were performed). Final $LEF_{cutoff}$ was obtained at $P$-value=0.10 from the distribution of LEF of permuted CRD. (B) Distribution of **(B)** Lead eigen factor of CRD called on $m_{peaks\_PEER} \times n_{samples}$ after filtering using $LEF_{cutoff}$ obtained from the previous step (A), stratified by three datasets H3K4me3 NeuN+, H3K27ac NeuN+ and H3K27ac Tissue. **(C)** Boxplot depicting the log of number of peaks in CRDs called on $m_{peaks\_PEER15} \times n_{samples}$ matrices of H3K4me3 NeuN+, H3K27ac NeuN+ and H3K27ac Tissue. **(D)** Distribution of proportion of enhancers (peaks that are located > ±3Kb from TSS) within the CRDs called on $m_{peaks\_PEER15} \times n_{samples}$ matrices of H3K4me3 NeuN+, H3K27ac NeuN+ and H3K27ac Tissue. **(E)** Proportions of CRDs containing only promoter-promoter, enhancer-enhancer, or a mixture of enhancer-promoter as a function of the number of peaks contained in CRDs across three groups. **(F)** Pairwise correlation of all histone peaks that are within the H3K4me3 NeuN+ (purple), H3K27ac NeuN+ (blue) and H3K27ac Tissue (green), and histone peaks that outside the CRDs (gray) as a function of distance. The x-axis is on a log scale. **(G)** Percentage of CRDs overlapping with TADs stratified by the number of overlapped TADs. **(H)** Fraction of TADs overlapping with CRDs stratified by the number of overlapped CRDs.

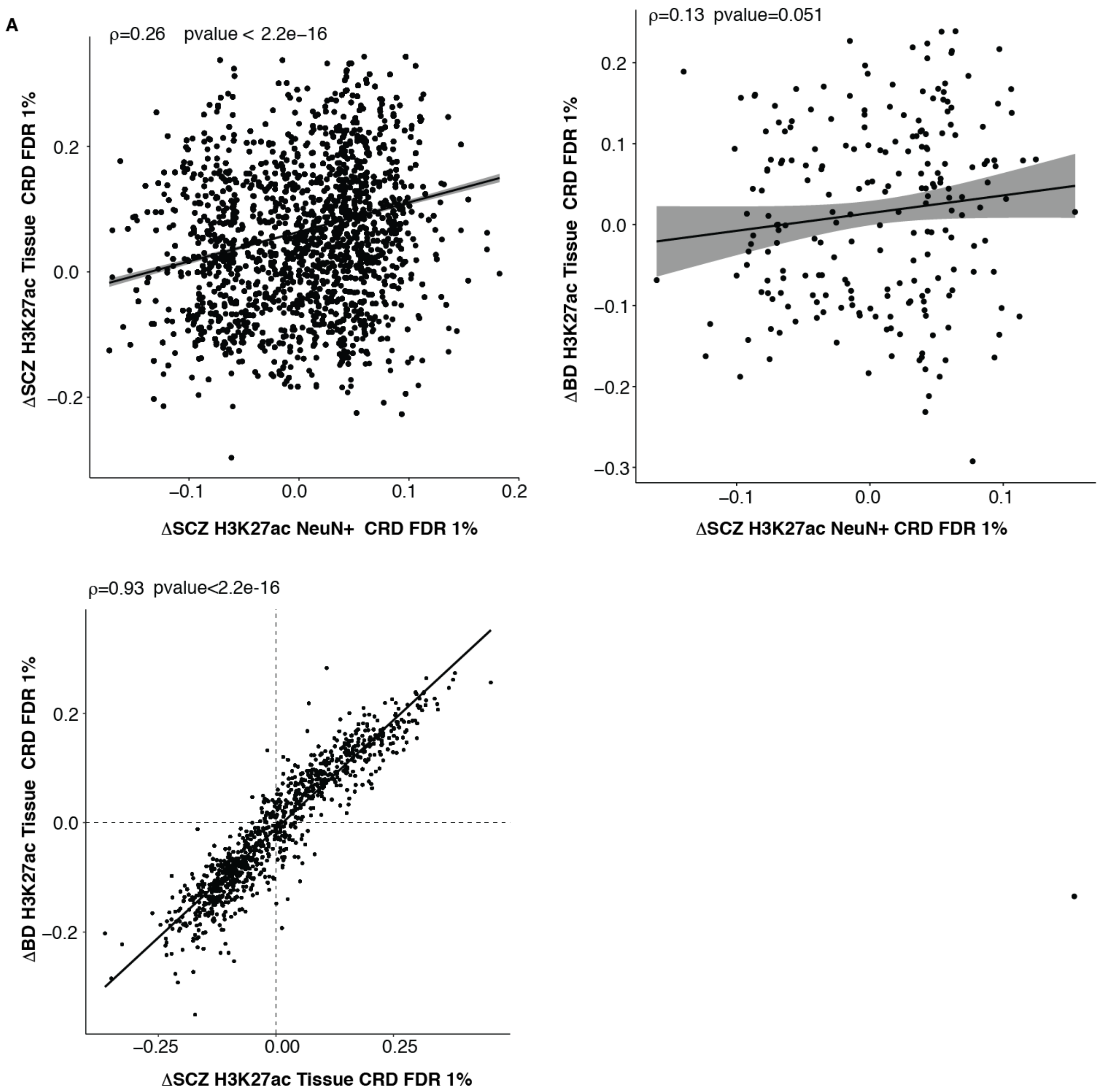


**Figure S10 | Concordance between sets of dysregulated CRDs across and within the studies. (A)** Left plot shows the spearman correlation between $log_2$FC (SCZ vs. controls) H3K27ac NeuN+ CRDs and $log_2$FC (SCZ vs. controls) H3K27ac Tissue at FDR 1% and right plot shows correlation between $log_2$FC (SCZ vs. controls) H3K27ac NeuN+ and $log_2$FC (BD vs. controls) H3K27ac Tissue at FDR 1% **(B)** Correlation between $log_2$FC (SCZ vs. controls) H3K27ac Tissue at FDR 5% and $log_2$FC (BD vs. controls) H3K27ac Tissue.

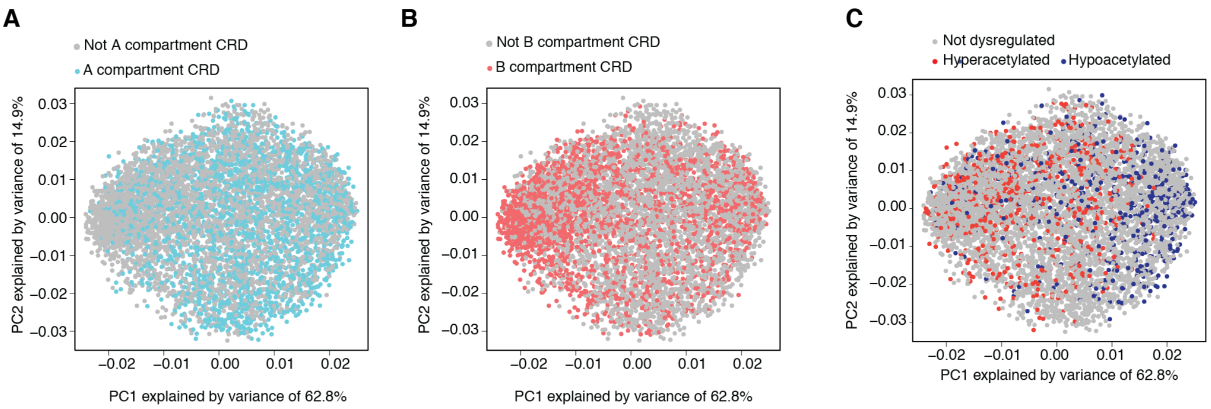


**Figure S11 | Demonstration of chromatin structure-activity relationship in CRDs.** Principal component analysis of H327ac NeuN+ CRD interaction map. Coordinates of the CRDs for Principal Component 1 (PC1) versus Principal Component 2 (PC2), together with the proportions of variance explained. Outcome of a PCA applied on annotated CRDs stratified by **(A)** A compartment CRDs in cyan **(B)** by B compartment CRDs in salmon color and all other CRDs in gray and **(C)** by dysregulation as hyperacetylated CRDs in red and hypoacetylated CRDs in blue

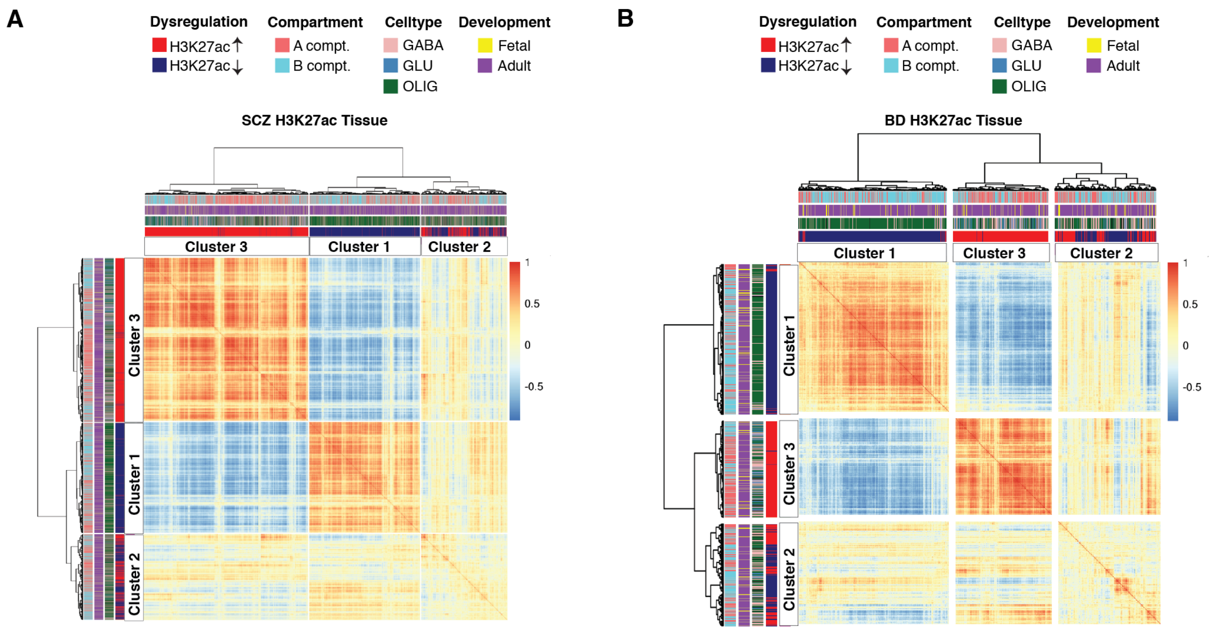


**Figure S12 | CRD interaction Map:** Classifiers for annotated CRDs are dysregulation (hypo vs hyper-acetylation in blue and red respectively), compartments (A compt, B compt and not annotated CRDs in salmon, cyan and gray respectively), cell type (GABA, GLU, OLIG in rose, blue and green respectively) and development (fetal, adult in yellow and purple respectively) (**A**) K-means clustering of CRD interaction map of SCZ-sensitive and **(B)** BD-sensitive H3K27ac Tissue CRDs into three large clusters; notice striking separation of cluster 1 representing hypoacetylated H3K27ac CRDs enriched for oligodendrocytes specific peaks, and clusters 2 and 3 overwhelmingly defined by hyperacetylated H3K27ac CRDs with GLU excitatory neuron-specific peak.

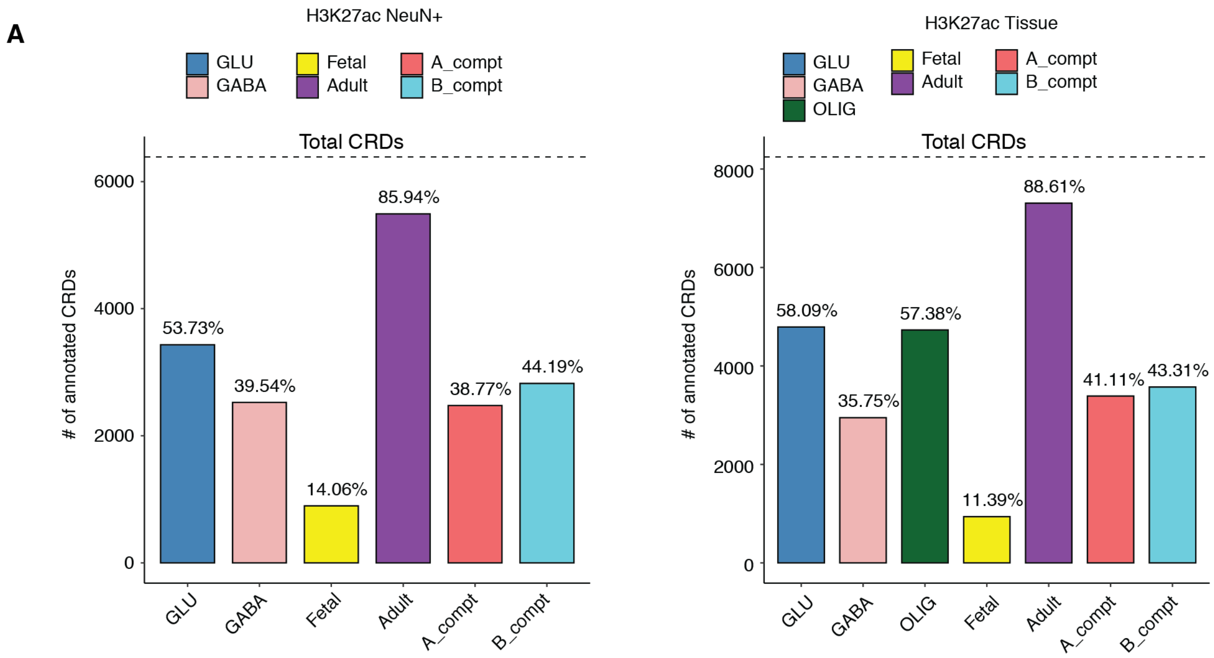


**Figure S13 | Annotated CRDs**. **(A)** Number of annotated CRDs to GLU (glutamatergic), GABA (gabaergic), fetal, adult, A compartment (A_compt) and B compartment (B_compt) of H3K27ac NeuN+ CRDs (right) and H3K27ac Tissue (left). The percentages above the bar plot are % of total CRDs annotated to a specific category. The dotted line shows the Total number of CRDs.

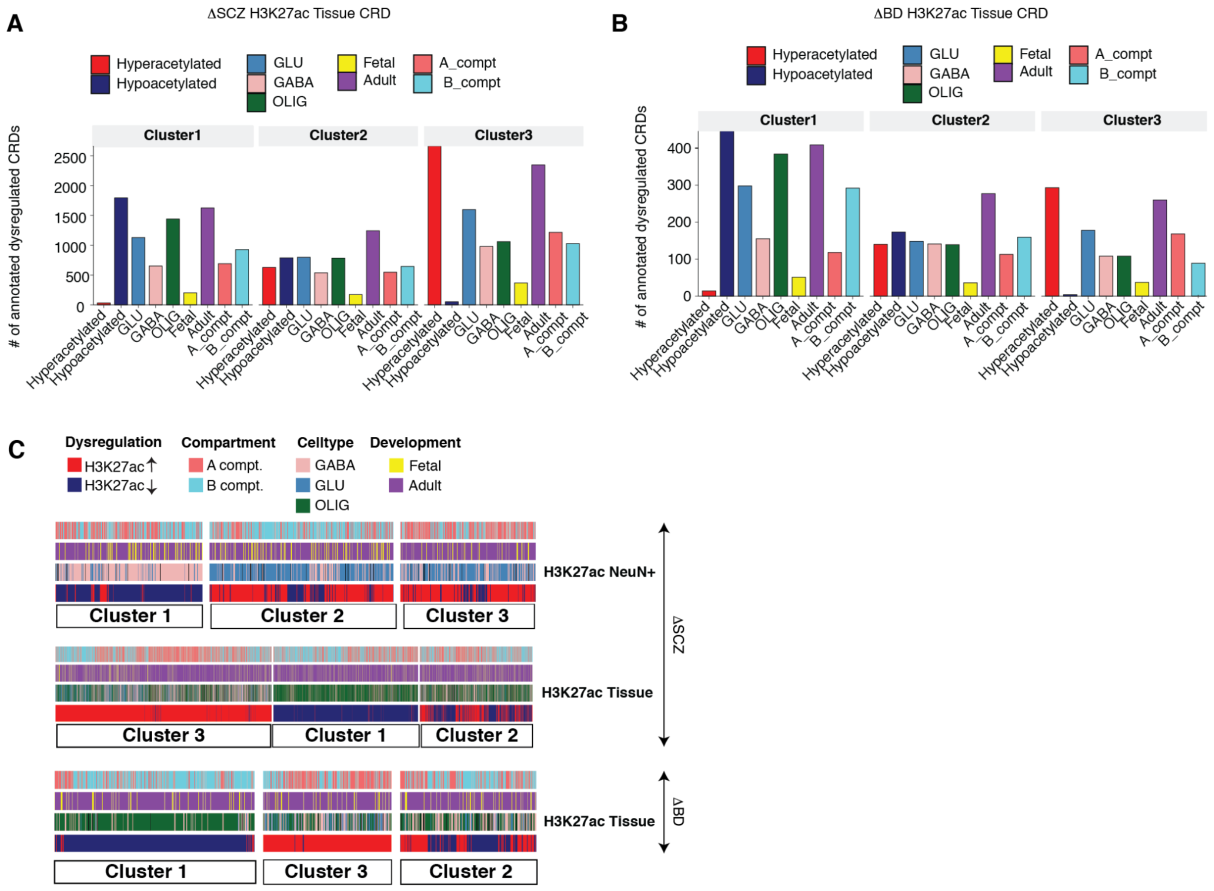


**Figure S14 | Clusters of disease specific CRD interaction map**. Classifiers for annotated disease specific CRDs are dysregulation (hypo vs hyper-acetylation), cell type (GABA, GLU, OLIG), development (fetal, adult), and compartments (A compt and B compt). Number of CRDs in every cluster stratified by every classifier for SCZ specific **(A)** and BD specific **(B)** analyses in H3K27ac Tissue. **(C)** Illustration of visual concordance of k-means clustering of CRD interaction map of SCZ-specific H3K27ac NeuN+ and H3K27ac Tissue and BD specific H3K27ac Tissue CRDs into three large clusters. Notice striking separation of cluster 1 representing hypoacetylated CRDs and clusters 2 and 3 overwhelmingly defined by hyperacetylated H3K27ac CRDs.

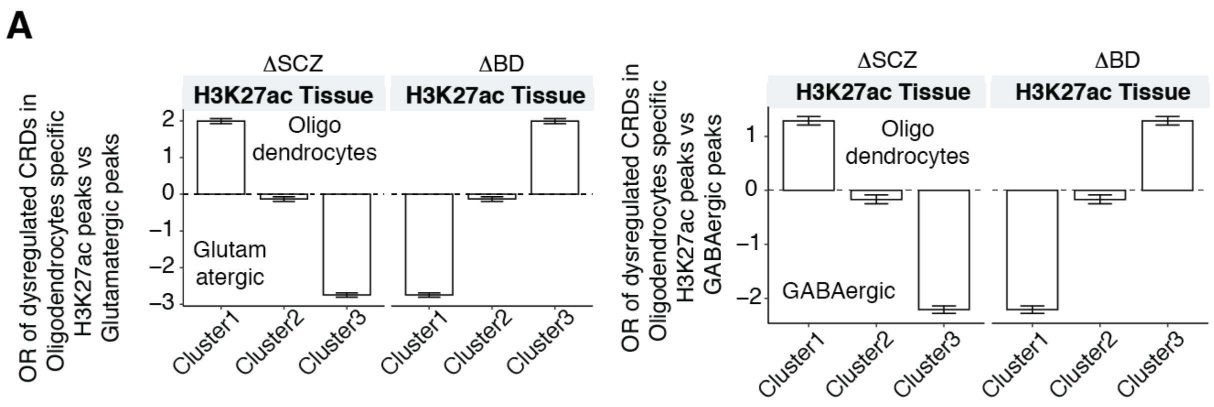


**Figure S15 | Enrichment of disease specific H3K27ac Tissue CRDs in oligodendrocytes. (A)** Odds ratio of SCZ and BD sensitive H3K27ac Tissue CRDs in oligodendrocytes vs glutamatergic specific H3K27ac peaks and **(B)** oligodendrocytes vs gabaergic specific H3K27ac peaks with genome wide CRDs as background stratified by cluster 1, 2 and 3.

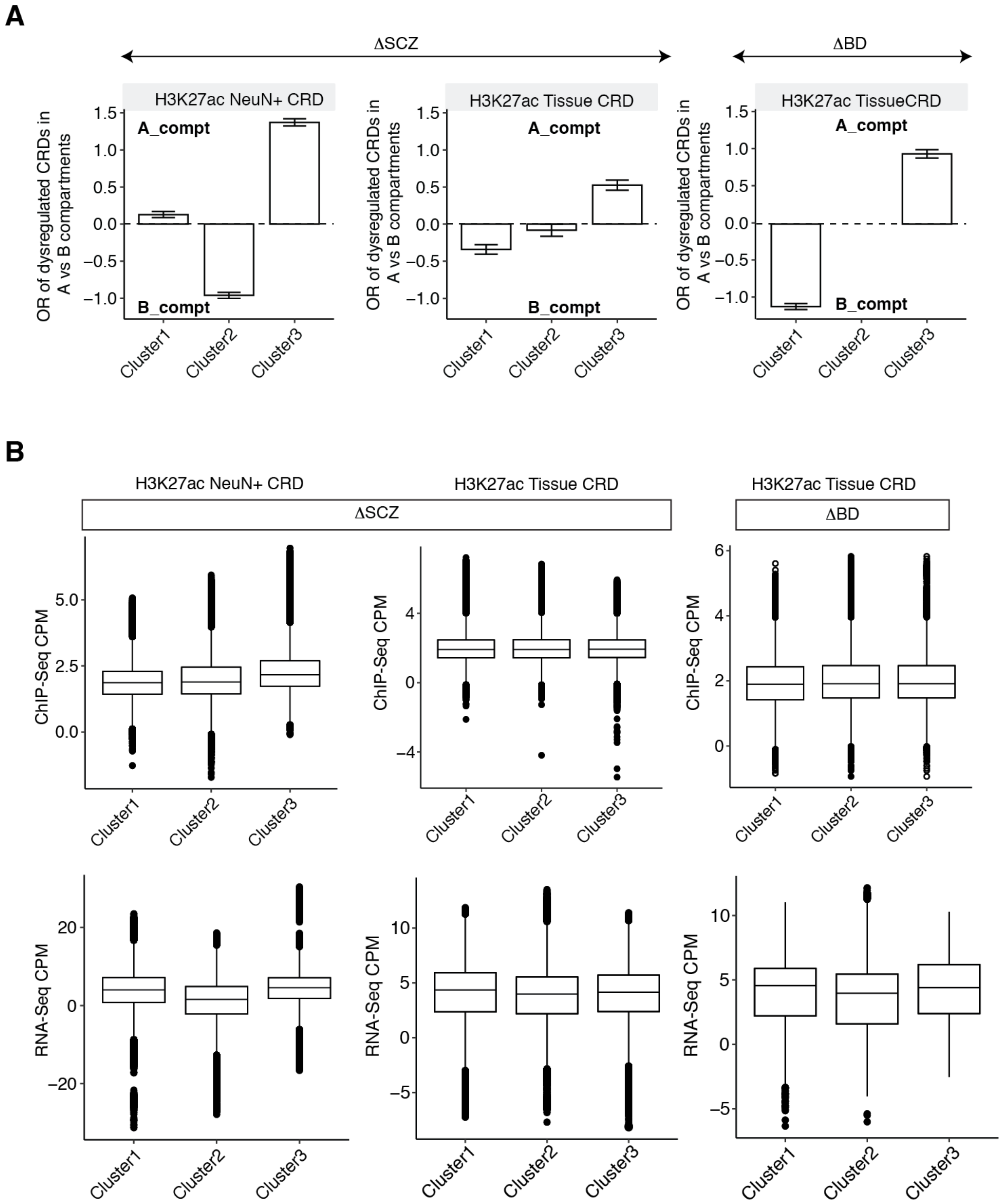


**Figure S16 | Spatial organization of disease specific CRDs. (A)** Odds ratio of SCZ sensitive H3K27ac NeuN+ and H3K27ac Tissue CRDs and BD sensitive H3K27ac Tissue CRDs in A vs B compartments from PFC HiC datasets with genome wide CRDs as background stratified by cluster 1, 2 and 3. **(B)** Box plot of ChIP-Seq expression of peaks and RNA-Seq expression of genes that are within SCZ sensitive H3K27ac NeuN+ and H3K27ac Tissue CRDs and BD sensitive H3K27ac Tissue CRDs stratified cluster 1,2 and 3.

A

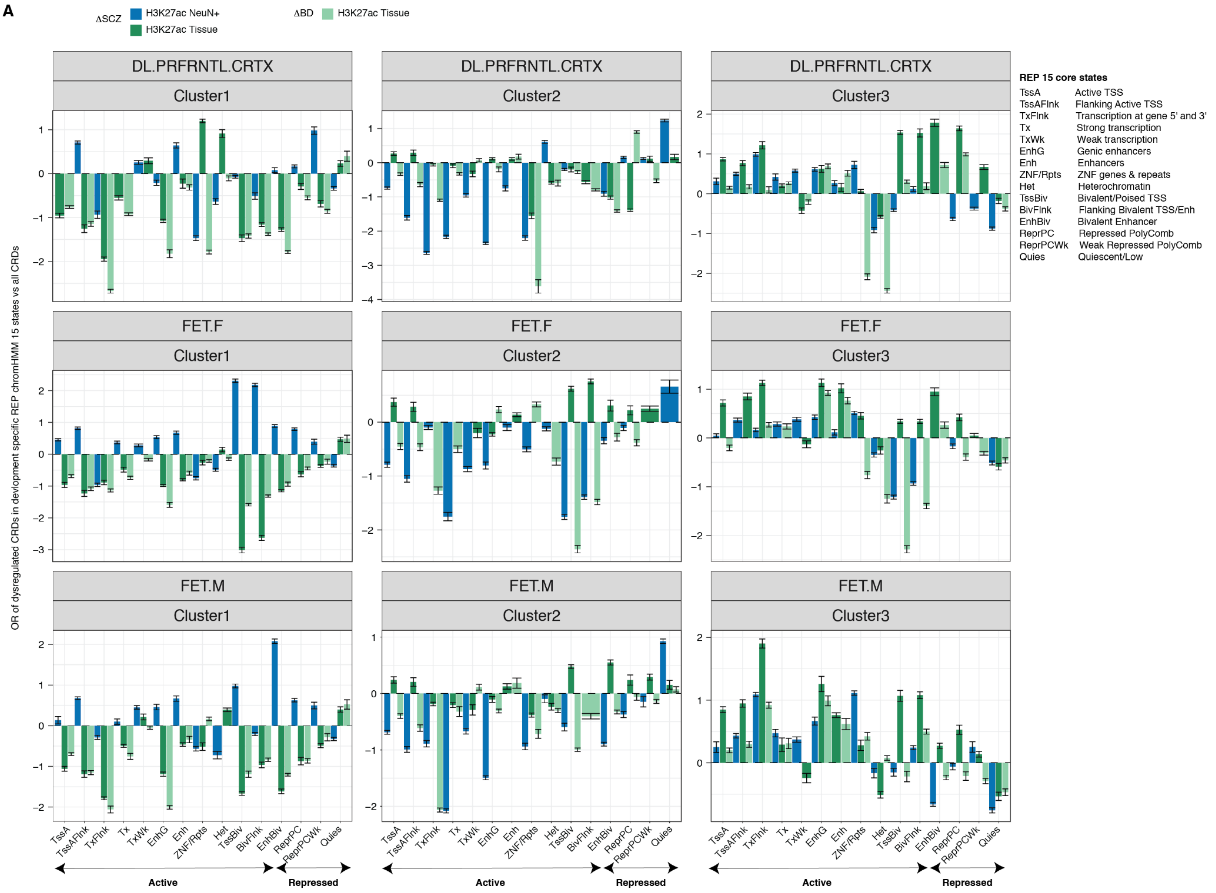


**Figure S17 | REP ChromHMM annotations of the DLPFC adult and fetal brain genome. (A)** Odds ratio of SCZ sensitive H3K27ac NeuN+ (blue) and H3K27ac Tissue (dark green) CRDs and BD sensitive H3K27ac Tissue (light green) CRDs in 15 chromHMM states from DLPFC adult brain(first row), fetal female brian(middle row) and fetal male brain (bottom row) with genome wide CRDs as background stratified by cluster 1, 2 and 3. X-axis abbreviations are explained in the left side of the figure.

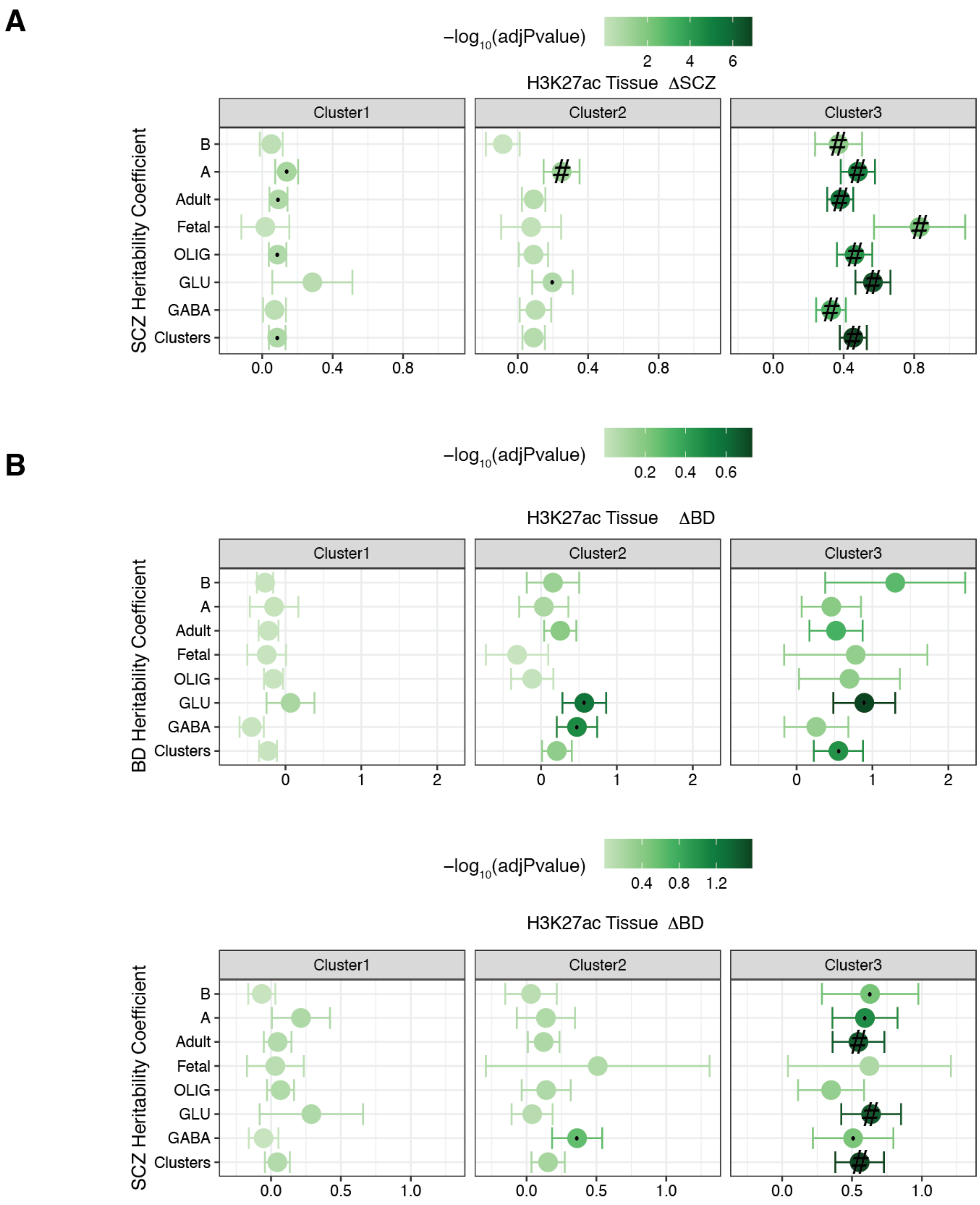


**Figure S18 | Cluster 3 is strongly enriched for SCZ and BD risk variants. (A)** SCZ heritability coefficients of genetic variants overlapping histone peaks from SCZ sensitive CRDs in H3K27ac Tissue stratified by cluster 1, 2 and 3. **(B)** BD (first row) and SCZ (second row) heritability coefficients of genetic variants overlapping histone peaks from dysregulated CRDs in BD sensitive H3K27ac tissue CRDs stratified by cluster 1, 2 and 3. The overlap of peaks with genetic variants was assessed using LD score regression. "#": Significant for enrichment in LD score regression after FDR correction of multiple testing across all tests in the plot (Benjamini & Hochberg); "·": Nominally significant for enrichment

A

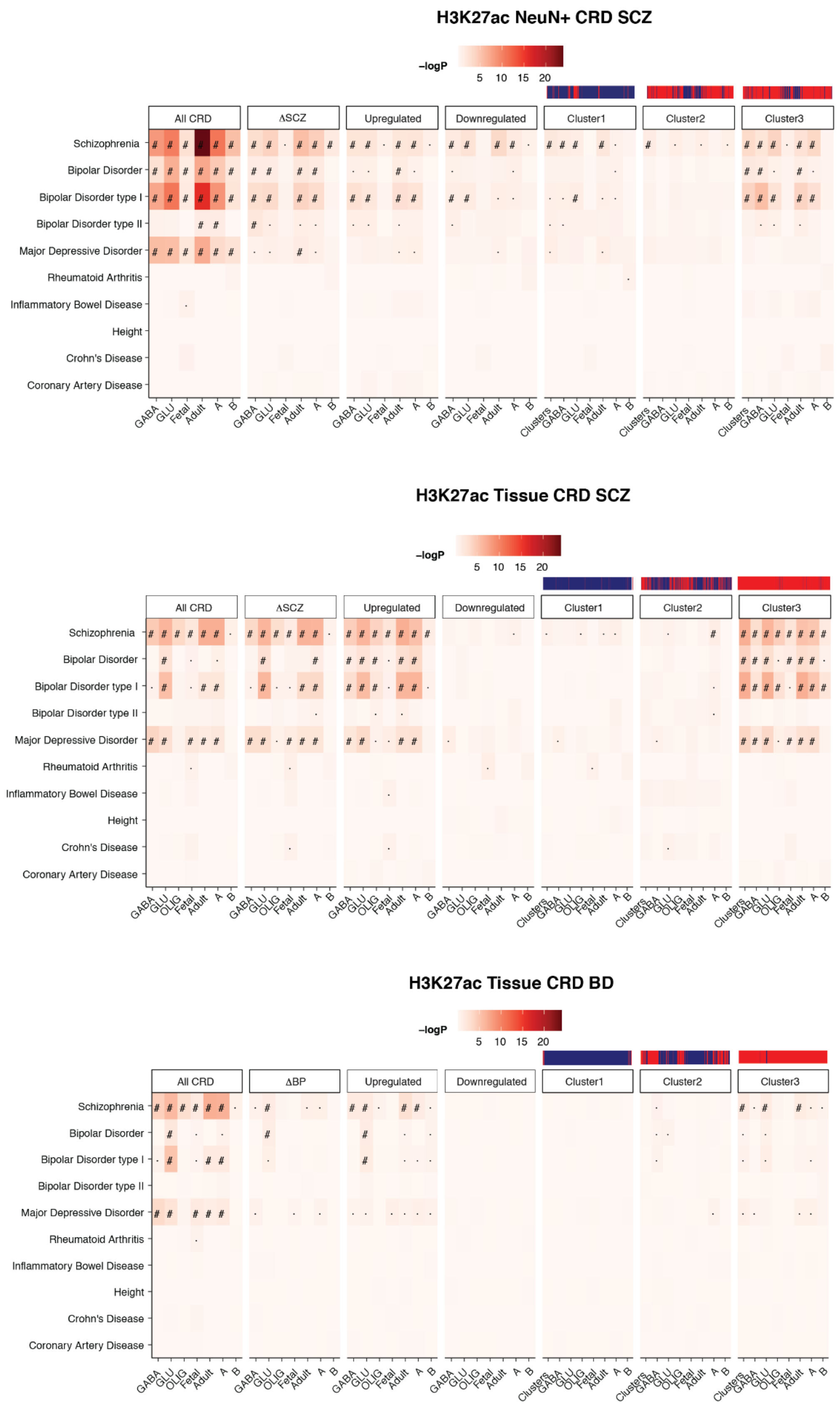


B

C

**Figure S19 | Cluster 3 is strongly enriched for risk variants associated with psychiatric traits.** Heatmap of enrichment *P*-values of brain and non-brain related GWAS traits in overlapping histone peaks within SCZ sensitive **(A)** H3K27ac NeuN+, **(B)** H3K27ac Tissue and **(C)** BD sensitive H3K27ac Tissue CRDs stratified cluster1, 2 and 3. The overlap of peaks with genetic variants was assessed using LD score regression. ”#”: Significant for enrichment in LD score regression after FDR correction of multiple testing across all tests in the plot (Benjamini & Hochberg); ”·”: Nominally significant for enrichment.

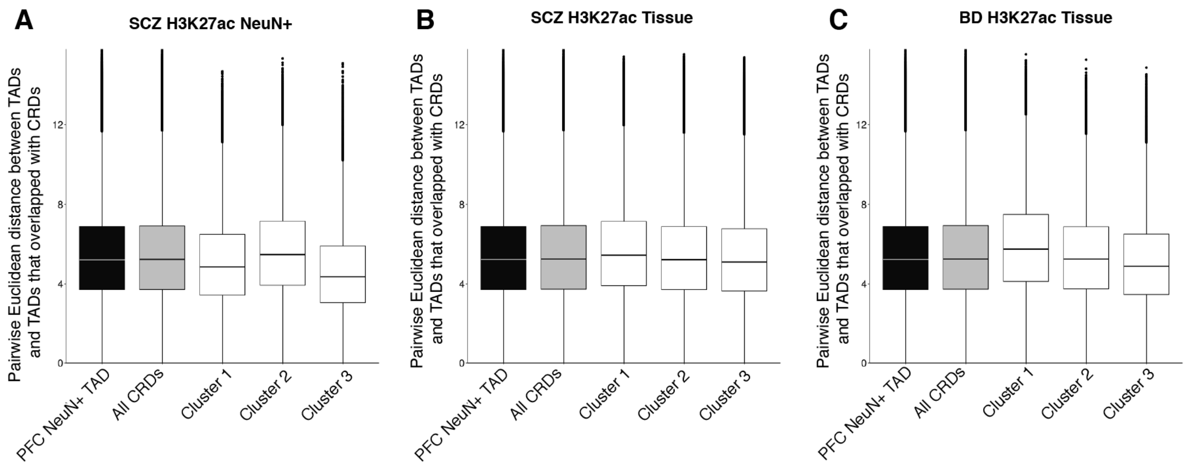


**Figure S20 | Three dimensional distance of diseased TADs in a three dimensional genome. (A)** Pairwise three dimensional distance between coordinates of all TADs (NeuN+ TAD), TADs overlapping with genome wide CRDs and dysregulated CRDs from cluster1, 2 and 3 from SCZ H3K27ac NeuN+ (left), SCZ H3K27ac Tissue (middle) and BD H3K27ac Tissue (right) .

#### Supplementary Tables

**Table S1**: Metadata of samples in study-1 and study-2.

**Table S2**: Genomic coordinates of consensus peaks.

**Table S3**: Differential analysis of peaks across SCZ cases and controls and across BD cases and controls.

**Table S4**: GWAS enrichment of brain traits and non-brain related traits in peaks stratified by differentially upregulated and downregulated and promoters and enhancers.

**Table S5**: Metadata of rat PFC brain samples and differential analysis of antipsychotics drugs.

**Table S6**: Genomic coordinates of identified CRDs.

**Table S7**: Annotation and differential analysis of CRDs.

**Table S8**: Genomic coordinates of H3K27ac Gabaergic, glutamatergic and oligodendrocytic specific peaks.

**Table S9**: Genomic coordinates of CRDs clusters in NeuN+ TAD 3D structure and pathway analysis of clustered NeuN+ TADs.

#### References (Methods)


1. Ghose, S., Gleason, K. A., Potts, B. W., Lewis-Amezcua, K. & Tamminga, C. A. Differential expression of metabotropic glutamate receptors 2 and 3 in schizophrenia: a mechanism for antipsychotic drug action? *Am. J. Psychiatry* **166**, 812–820 (2009).
2. Gao, X.-M., Cooper, T., Suckow, R. F. & Tamminga, C. A. Multidose risperidone treatment evaluated in a rodent model of tardive dyskinesia. *Neuropsychopharmacology* **31**, 1864–1868 (2006).
3. Wang, D. *et al.* Comprehensive functional genomic resource and integrative model for the human brain. *Science* **362**, (2018).
4. Kundakovic, M. *et al.* Practical Guidelines for High-Resolution Epigenomic Profiling of Nucleosomal Histones in Postmortem Human Brain Tissue. *Biol. Psychiatry* **81**, 162–170 (2017).
5. Jiang, Y., Matevossian, A., Huang, H.-S., Straubhaar, J. & Akbarian, S. Isolation of neuronal chromatin from brain tissue. *BMC Neurosci.* **9**, 42 (2008).
6. Bolger, A. M., Lohse, M. & Usadel, B. Trimmomatic: a flexible trimmer for Illumina sequence data. *Bioinformatics* **30**, 2114–2120 (2014).
7. Li, H. & Durbin, R. Fast and accurate short read alignment with Burrows-Wheeler transform. *Bioinformatics* **25**, 1754–1760 (2009).
8. Landt, S. G. *et al.* ChIP-seq guidelines and practices of the ENCODE and modENCODE consortia. *Genome Res.* **22**, 1813–1831 (2012).
9. Fort, A. *et al.* MBV: a method to solve sample mislabeling and detect technical bias in large combined genotype and sequencing assay datasets. *Bioinformatics* **33**, 1895–1897 (2017).
10. Zhang, Y. *et al.* Model-based analysis of ChIP-Seq (MACS). *Genome Biol.* **9**, R137 (2008).
11. Amemiya, H. M., Kundaje, A. & Boyle, A. P. The ENCODE blacklist: identification of problematic regions of the genome. *Sci. Rep.* **9**, 9354 (2019).
12. Liao, Y., Smyth, G. K. & Shi, W. featureCounts: an efficient general purpose program for assigning sequence reads to genomic features. *Bioinformatics* **30**, 923–930 (2014).
13. Robinson, M. D., McCarthy, D. J. & Smyth, G. K. edgeR: a Bioconductor package for differential expression analysis of digital gene expression data. *Bioinformatics* **26**, 139–140 (2010).
14. Hunt, G. J., Freytag, S., Bahlo, M. & Gagnon-Bartsch, J. A. dtangle: accurate and robust cell type deconvolution. *Bioinformatics* **35**, 2093–2099 (2019).
15. Neath, A. A. & Cavanaugh, J. E. The Bayesian information criterion: background, derivation, and applications. *WIREs Comp Stat* **4**, 199–203 (2012).
16. Yu, G., Wang, L.-G. & He, Q.-Y. ChIPseeker: an R/Bioconductor package for ChIP peak annotation, comparison and visualization. *Bioinformatics* **31**, 2382–2383 (2015).
17. Ernst, J. & Kellis, M. Chromatin-state discovery and genome annotation with ChromHMM. *Nat. Protoc.* **12**, 2478–2492 (2017).
18. Girdhar, K. *et al.* Cell-specific histone modification maps in the human frontal lobe link schizophrenia risk to the neuronal epigenome. *Nat. Neurosci.* **21**, 1126–1136 (2018).
19. Ritchie, M. E. *et al.* limma powers differential expression analyses for RNA-sequencing and microarray studies. *Nucleic Acids Res.* **43**, e47 (2015).
20. Viechtbauer, W. Conducting Meta-Analyses in R with the metafor Package. *J. Stat. Softw.* **36**, (2010).
21. McLean, C. Y. *et al.* GREAT improves functional interpretation of cis-regulatory regions. *Nat. Biotechnol.* **28**, 495–501 (2010).
22. Finucane, H. K. *et al.* Partitioning heritability by functional annotation using genome-wide

association summary statistics. *Nat. Genet.* **47**, 1228–1235 (2015).
23. Stegle, O., Parts, L., Piipari, M., Winn, J. & Durbin, R. Using probabilistic estimation of expression residuals (PEER) to obtain increased power and interpretability of gene expression analyses. *Nat. Protoc.* **7**, 500–507 (2012).
24. Hoffman, G. E., Bendl, J., Girdhar, K. & Roussos, P. decorate: differential epigenetic correlation test. *Bioinformatics* **36**, 2856–2861 (2020).
25. ENCODE Project Consortium. An integrated encyclopedia of DNA elements in the human genome. *Nature* **489**, 57–74 (2012).
26. Bendl, J. *et al.* The three-dimensional landscape of chromatin accessibility in Alzheimer's disease. *BioRxiv* (2021) doi:10.1101/2021.01.11.426303.
27. Kozlenkov, A. *et al.* A unique role for DNA (hydroxy)methylation in epigenetic regulation of human inhibitory neurons. *Sci. Adv.* **4**, eaau6190 (2018).
28. Kumar, V. *et al.* Uniform, optimal signal processing of mapped deep-sequencing data. *Nat. Biotechnol.* **31**, 615–622 (2013).
29. Li, M. *et al.* Integrative functional genomic analysis of human brain development and neuropsychiatric risks. *Science* **362**, (2018).
30. Kozlenkov, A. *et al.* Substantial DNA methylation differences between two major neuronal subtypes in human brain. *Nucleic Acids Res.* **44**, 2593–2612 (2016).
31. Forgy, E. Cluster analysis of multivariate data : efficiency versus interpretability of classifications. *undefined* (1965).
32. Servant, N. *et al.* HiC-Pro: an optimized and flexible pipeline for Hi-C data processing. *Genome Biol.* **16**, 259 (2015).
33. Ramírez, F. *et al.* High-resolution TADs reveal DNA sequences underlying genome organization in flies. *Nat. Commun.* **9**, 189 (2018).
34. Abdennur, N. & Mirny, L. A. Cooler: scalable storage for Hi-C data and other genomically labeled arrays. *Bioinformatics* **36**, 311–316 (2020).
35. Paulsen, J. *et al.* Chrom3D: three-dimensional genome modeling from Hi-C and nuclear lamin-genome contacts. *Genome Biol.* **18**, 21 (2017).
36. Paulsen, J., Liyakat Ali, T. M. & Collas, P. Computational 3D genome modeling using Chrom3D. *Nat. Protoc.* **13**, 1137–1152 (2018).
